# Body stoichiometry of heterotrophs: assessing drivers of interspecific variations in elemental composition

**DOI:** 10.1101/2020.04.08.027656

**Authors:** Benjamin Andrieux, Juliette Signor, Vincent Guillou, Michael Danger, Franck Jabot

## Abstract

**Aim:** To document how body stoichiometry of heterotrophs varies globally and to assess phylogenetic, trophic, habitat and body mass drivers of this interspecific variation in elemental composition, focusing on carbon (C), nitrogen (N) and phosphorus (P).

**Location:** Worldwide.

**Time period:** 1930 – 2018.

**Major taxa studied:** Amphibians, fishes (Euteleosteomorpha and Otomorpha), invertebrates, mammals, microbes and sauropsids (birds and reptiles).

**Methods:** We compiled from the scientific literature a global database of body elemental composition of heterotrophs in marine, freshwater and terrestrial realms. We used model selection and ANCOVA analyses to investigate the proportion of variance in elemental composition explained by taxonomic groups, diet, habitat and body mass. We assessed the phylogenetic signal in body stoichiometry using Blomberg’s K and Pagel’s λ statistics. We assessed the phylogenetic structure of interspecific variation in body stoichiometry using mixed models with nested taxonomic levels as random factors. We finally assessed the co-variation in elemental composition using linear models.

**Results:** Our database currently gathers 17848 independent observations on 1491 species. Body elemental composition was found to be widely variable among species with the four assessed drivers significantly contributing to this variation. Taxonomic group is the strongest contributor to interspecific variance for the stoichiometric traits studied, followed by habitat, diet and body mass. More precisely, stoichiometric traits are generally variable at the three taxonomic levels studied (class, order and family), resulting in a significant but relatively modest phylogenetic signal. Finally, we found significant co-variation among the three body elemental contents, resulting in taxonomic group-specific C:N:P spectrums.

**Main conclusions:** Our global synthesis of body stoichiometry of heterotrophs reveals a strong interspecific variability that is only modestly explained by the species attributes investigated: body mass, habitat and diet. It further reveals that this taxonomically structured residual variation in body stoichiometry seems to be constrained along taxonomic group-specific elemental spectrums.

## INTRODUCTION

Ecological stoichiometry theory revolves around the main idea that the growth and reproduction of an organism depends on its ability to acquire nutrients in appropriate proportions from its environment (Sterner & Elser, 2002). Following this theory, the body elemental composition of an organism is thus a critical parameter of its ecology. Autotrophic life forms generally harbor a substantial plasticity in terms of elemental composition that greatly varies in response to nutrient and light modification of their environment (Aerts & Chapin, 1999; Duarte, 1992). In contrast, heterotrophic organisms generally present much less plastic variation in their elemental composition (Lemmen et al., 2019), a phenomenon called stoichiometric homeostasis. Although the idea of a strict homeostasis of heterotrophs has been recurrently falsified (Persson et al., 2010), it constitutes a convenient approximation, at least for metazoans. This approximation is further used in many stoichiometric models of community and ecosystem dynamics (Andersen et al., 2004).

Understanding how body nutrient contents vary among organisms and assessing putative drivers of such variations constitute a cornerstone of ecological stoichiometry. Various hypothesized drivers of stoichiometric variations among organisms have been put forward. The growth rate hypothesis (Elser et al., 2003; Elser, Dobberfuhl, MacKay, & Schampel, 1996) states that fast growing organisms should have lower N:P ratios than slow growing ones. Besides, organisms with large metabolic rates may consume larger amounts of C and thus may have lower C:N and C:P ratios. Since larger organisms tend to have lower metabolic rates per unit mass (Brown, Gillooly, Allen, Savage, & West, 2004) and lower growth rates, body mass should be positively related to C:N, C:P and N:P. Diet of organisms may also influence their stoichiometric needs. Since predators have access to preys with larger N and P contents, they may have been selected (or less counter-selected) to have larger N and P contents (Sterner & Elser, 2002). Finally, some phylogenetically-conserved traits may profoundly influence stoichiometric constraints of organisms. For instance, the presence of a skeleton should increase P content (Simkiss & Wilbur, 1989), while homeothermic organisms are likely to have larger C needs than poikilothermic individuals (Sterner & Elser, 2002).

Many works have investigated stoichiometric patterns in specific taxonomic groups (see our compilation in Table S1). To sum up these pieces of work, (i) most studies have focused on carbon (C), nitrogen (N) and phosphorus (P) contents of organisms, or on their associated ratios C:N, C:P and N:P; (ii) the putative drivers of elemental composition mainly focused on phylogeny/taxonomy, diet, body mass, ontogeny and habitat; (iii) empirical patterns and their magnitude are variable between the different taxonomic groups studied. More specifically, evolutionary history is a significant predictor of stoichiometric traits in various groups when analyzed with analyses of variances based on taxonomic groups, especially at large taxonomic levels (phyla, classes, orders): in terrestrial insects (Fagan et al., 2002; Martinson et al., 2008; Wiesenborn, 2013; Woods, Fagan, Elser, & Harrison, 2004), fishes (Hendrixson, Sterner, & Kay, 2007; McIntyre & Flecker, 2010; Vanni, Flecker, Hood, & Headworth, 2002), aquatic invertebrates (Evans-White, Stelzer, & Lamberti, 2005; González et al., 2018), lizards (González, Fariña, Kay, Pinto, & Marquet, 2011) and fungi (Zhang & Elser, 2017). In contrast, most of these studies found little or no phylogenetic signal in body stoichiometry, probably because of too low sample sizes (Blomberg, Garland, & Ives, 2003). Diet is also a recurrent predictor of body stoichiometry across taxonomic groups. Roughly, predators tend to have larger N and/or P contents than herbivores and/or detritivores in terrestrial insects (Fagan et al., 2002; González et al., 2011; Wiesenborn, 2013), fishes (McIntyre & Flecker, 2010) and aquatic invertebrates (Evans-White et al., 2005). In contrast with these general findings, Vanni et al. (2002) evidenced in their study similar P contents between predator and herbivorous fishes, and González et al. (2008) evidenced lower P contents in predators compared to detritivores in aquatic invertebrates. Finally, Martinson et al. (2008) evidenced similar N contents between herbivores and detritivores in terrestrial arthropods. Body mass is a less straightforward predictor of body stoichiometry. When studied at the intraspecific level, correlations between body mass and elemental contents are strongly species-specific in sign, significance and magnitude (Arbačiauskas, Lesutienė, & Gasiūnaitė, 2013; Hendrixson et al., 2007; but see Gonzàlez et al., 2018; Woods, 2004). When studied at the interspecific level, correlations between body mass and elemental contents are also variable across taxonomic groups with positive, negative, hump-shape or non-significant relationships. Moreover, allometric relationships tend to vanish when controlling for phylogenetic effects (e.g., González et al., 2018; González et al., 2011; Hendrixson et al., 2007). A fewer number of works have studied the effect of ontogeny on body stoichiometry. Fagan et al. (2002) did not find any ontogenetic signal in insects, while González et al. (2011) found larger C:N ratios in larvae than adult arthropods. Many studies failed to detect a significant effect of habitat on body stoichiometry or detected either a significant but low habitat effect or a species-specific habitat effect (Arbačiauskas et al., 2013 for crustaceans; Evans-White et al., 2005 for aquatic invertebrates; González et al., 2011 for arthropods and lizards; Milanovich, Maerz, & Rosemond, 2015 for salamanders). Some works however did succeed in evidencing habitat effects in fishes (Hendrixson et al., 2007), aquatic invertebrates (González et al., 2018) and fungi (Zhang & Elser, 2017).

As illustrated above, syntheses on the ecological stoichiometry of heterotrophs are until now taxonomically segmented with heterogeneous data analysis approaches and variable sets of stoichiometric drivers studied. They end up with contrasted or sometimes contradictory results that hamper a general understanding of the ecological stoichiometry of heterotrophs. The aim of the present study is thus to compile a unified database of heterotroph body elemental contents (C, N and P) and associated ratios (C:N, C:P and N:P), in order to conduct a general analysis across taxonomic groups. We will use this database to assess i) the variability of body elemental contents and their co-variations documented worldwide across heterotrophs; and ii) the importance of four putative drivers for these variations: evolutionary history, diet, body mass and habitat.

## METHODS

### Data compilation

We assembled three already compiled extensive datasets synthesizing information about aquatic animals (Vanni et al., 2017), fungi (Zhang & Elser, 2017) and insects (Fagan et al., 2002). We completed this dataset using published literature up to the end of August 2019. We conducted this literature search on ISI Web of Knowledge and Google Scholar. Search terms included ‘nutrient content’, ‘body compo*’, ‘nutrient compo*’, ‘chemical compo*’, ‘protein content’, carbon content’, nitrogen content’, ‘phosphorus content’ and ‘ecological stoichiometr*’ together with ‘ontogen*’, ‘allometr*’, ‘body size’, ‘trophic’, ‘trophy’, ‘diet*’. Preliminary attempts based on more specific keywords did not enable us to collect much data. We therefore relied on such broad keywords search with the limitation that we could not systematically consult all the articles found. We therefore do not pretend to have been exhaustive. In this literature search, we paid particular attention to collect data on taxa that were less informed by previous syntheses, especially mammals and sauropsids. To do this, we reiterated the search including group names and synonyms in the query. A list of the data sources is found in Appendix 1.

We included in our database all data with at least one observation of body carbon, nitrogen or phosphorus content or one of their ratio, except that we removed the values of C content for the fungi group when C content was virtually set to an average value (Zhang & Elser, 2017). We used the program ‘Plot Digitizer’ v2.6.8 to extract data from graphs. When only body protein content was given in a publication, we divided this number by 6.25 to estimate nitrogen content (Persson et al., 2010). We broadly taxonomically grouped organisms as amphibians, fishes (separating two distinct groups: Euteleosteomorpha and Otomorpha), invertebrates, mammals, microbes or sauropsids (birds and reptiles). We recorded the main diet of the organism as being either herbivore, carnivore, detritivore or omnivore. We recorded the main habitat of the organism as being either terrestrial, freshwater or marine. We finally used the log-transformed dry mass of an organism in subsequent statistical analyses. When necessary, we converted fresh (wet) mass to dry mass considering dry mass to be 28.82% of wet mass for mammals, 34.98% for birds, 26.80% for reptiles, 23.50% for fishes and 20.46% for amphibians. We obtained these ratios from the weighted average by number of observations for mammals (Arnould, Boyd, & Speakman, 1996; Gerhart, White, Cameron, & Russell, 1996; Kremen et al., 2013; Reilly & Fedak, 1990; Schlesinger & Potter, 1974; Spray & Widdowson, 1950; Studier, Sevick, & Wilson, 1994), birds (Grimshaw, Ovington, Betts, & Gibb, 1958; Kremen et al., 2013; Sturges, Holmes, & Likens, 1974), reptiles (Barron, 1997; Dierenfeld, Alcorn, & Jacobsen, 2002), fishes (Lantry & O’Gorman, 2007) and amphibians (Dierenfeld et al., 2002; MacCracken & Stebbings, 2012; Takahara, Miyasaka, Genkai-Kato, & Kohmatsu, 2008). When diet, body mass or habitat of the organism were not mentioned in the publication, we gathered them from other publications and specified the corresponding references in the database. Typing errors for species names were checked by using the Open Tree Taxonomy’s taxon resolution name service together with the ‘rotl’ R package(Michonneau, Brown, Winter, & Fitzjohn, 2016), from which we also imputed the taxonomy levels for most of the species. We also assembled a dated phylogenetic tree for a subset (n = 344) of the species present in the dataset with the ‘datelife’ R package (Sánchez-Reyes & O’Meara, 2019).

### Elemental content variability

We assessed the range of elemental contents for each taxonomic group. We also assessed the presence of “stoichiometric spectrums”, that is of significant co-variations between body C, N and P contents among heterotrophs. We visually assessed these spectrums using three-dimensional plots. We then statistically tested the significance of these spectrums using multiple regressions explaining C content as a function of N and P contents both on the full dataset and on each taxonomic group separately. We also assessed the three pairwise Pearson’s correlations between C, N and P contents both on the full dataset and on each taxonomic group separately.

### Stoichiometric drivers

We first assessed the significance of four stoichiometric drivers (taxonomic group, diet, body mass and habitat) on elemental contents (C, N and P) and elemental ratios (C:N, C:P, N:P) with additive linear models without interactions between drivers. We built 15 models corresponding to all the possible combinations of these four drivers and performed a model selection based on the Akaike information criterion (AIC). We retained the ‘best’ model (i.e., the model that best fitted the data) with the lowest AIC plus the other models with a difference in AIC below 10 (ΔAIC < 10, Burnham & Anderson, 2002). We also computed the corresponding model weights (W). We then assessed the magnitude of these drivers using partial eta-squared (partial η^2^) on the full model (with the four drivers). Partial η^2^ compute the ratio between the variance explained by a stoichiometric driver and the sum of this variance plus the variance of the residuals. It therefore measures the proportion of the remaining variance explained by a driver once the effects of the other drivers are factored out.

For visualization purposes, we also used the ‘sequential ANCOVA’ procedure described in Fagan et al. (2002). This consists, for each driver in turn, in fitting an additive linear model with the three other drivers, and in assessing the effect of the focal driver on the residuals of this model. By doing this, we are conservative on the effect of each driver. For categorial variables, comparison among groups was done with non-parametric Kruskal-Wallis rank sum test with post-hoc Dunn’s test of multiple comparisons with Bonferroni corrections. Before analyses, body mass was log-transformed. The body mass of microbes was not informed in the studied data sets, so this group was not included in these analyses.

### Evolutionary history

We computed the phylogenetic signal of stoichiometric traits (C, N, P, C:N, C:P, N:P) with two statistics: Blomberg’s K (Blomberg et al., 2003) and Pagel’s λ (Pagel, 1999). The significance of phylogenetic signal was also assessed with both the randomization test of Blomberg et al. (2003) and the likelihood ratio test associated with Pagel’s λ. For such phylogenetic analyses, we used species-averaged values of stoichiometric traits. We also fitted linear mixed models for each stoichiometric trait separately, using diet, habitat and body mass as fixed factors and taxonomic classes, orders and families as nested random factors using the ‘nlme’ R package (Pinheiro, Bates, DebRoy, Sarkar, & R Core Team, 2020). The variance in stoichiometric traits explained by the fixed factors was evaluated with the marginal R^2^ and the variance explained both by the fixed and random factors by the conditional R^2^ using the ‘MuMIn’ R package (Barton, 2019). In addition, we assessed the variance components associated with each taxonomic level by using the ‘ape’ R package (Paradis & Schliep, 2018).

All statistical analyses were performed with R v3.6.1 (R development core team 2017). We used the package ‘AICcmodavg’ for model selection (Mazerolle, 2019) and ‘yarrr’ (Phillips, 2017) and ‘plot3D’ (Soetaert, 2019) for data visualization. ANCOVAs (type III) were performed using the ‘car’ package (Fow & Weisberg, 2019) together with the ‘FSA’ (Ogle, Wheeler, & Dinno, 2020) and ‘rcompanion’ (Mangiafico, 2020) R packages for Dunn’s tests and compact letter display for lists of comparisons. We managed phylogenetic data with ‘ape’ package (Paradis & Schliep, 2018) and phylogenetic signal was calculated with the ‘phytools’ package (Revell, 2012).

## RESULTS

### Data compilation

Our database gathers 56 sources of information (original publications or previous syntheses). It compiles information of 373 studies and contains 17,848 independent stoichiometric observations on 1,491 species. It successfully compiles information for the seven main taxonomic groups considered (amphibians, fishes-Euteleosteomorpha, fishes-Otomorpha, invertebrates, mammals, microbes and sauropsids), although with variable sample sizes (Fig. S2a). A diverse array of diets (Fig. S2b) and habitats (Fig. S2c) are sampled in most taxonomic groups. The range of body mass within each group is substantial (Fig. S2d). This compilation therefore enables us to conduct the statistical analyses envisioned, although we must recognize the fact that our sample of heterotrophs is substantially unbalanced among taxonomic groups, diets, habitats and body masses, and is factorially incomplete as inherent to such literature syntheses.

### Elemental content variability

Across the full dataset, elemental contents present limited variabilities, with C content (first and third quartiles at 41 and 48.1 respectively) varying less than N content (8.2;11) and P content (0.7;2.1). Still, some organisms present important deviations from these median ranges with C content varying from 4.1 to 65.3, N content from 0.2 to 19 and P content from 0.04 to 11.4. Within each taxonomic group studied, empirical variability in elemental contents is of similar magnitude, with largely overlapping distributions between the various groups (Fig. 1). Some differences between taxonomic groups are however noticeable: amphibians have above-average N contents (Fig. 1b), mammals have above-average C and N contents (Fig. 1a,b), microbes have below-average N and P contents (Fig. 1b,c), sauropsids have below-average C contents (Fig. 1a) and invertebrates have below-average P contents (Fig. 1c). Some invertebrate species present low to very low C and N contents, these species correspond to some marine gelatinous organisms. Also, there is less overlap in P contents between the different taxonomic groups (Fig. 1c). As a consequence of low body N and P content, microbes have larger C:N and C:P ratios than other organisms (Fig. 1d,e). Also, invertebrates have above-average N:P ratio (Fig. 1f) and sauropsids below-average C:P and N:P ratios (Fig. 1e,f).

**Figure 1.**
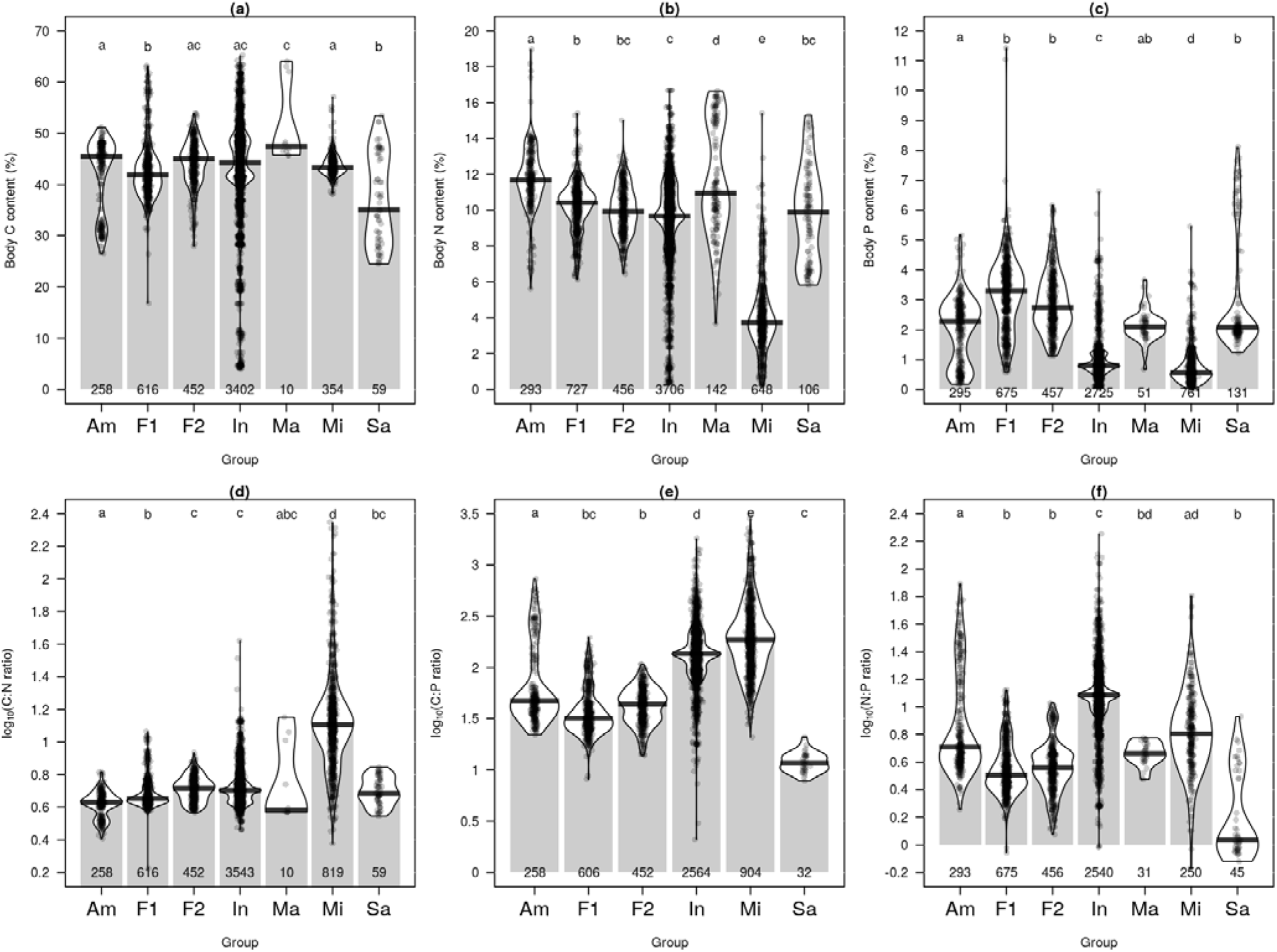
Elemental content variation among taxonomical groups. (a) Body C content. (b) Body N content. (c) Body P content. (d) Log-transformed C:N ratio. (e) Log-transformed C:P ratio. (f) Log-transformed N:P ratio. Am: Amphibians, F1: Fishes – clade Euteleosteomorpha, F2: Fishes – clade Otomorpha, In: Invertebrates, Ma: Mammals, Mi: Microbes, Sa: Sauropsids (birds and reptiles). Raw data (points) and distributions (kernel densities) are depicted with the number of observations (at the bottom for each panel). Different letters indicate significant differences with post-hoc Dunn’s tests of multiple comparisons with Bonferroni corrections that were performed before log-transformation.

We find a substantial amount of pairwise co-variation between elemental contents: when all taxonomic groups are analyzed together, there is a positive Pearson’s correlation coefficient of 0.43 between C and N contents, of 0.20 between N and P contents and a slight negative correlation of −0.05 between C and P contents. We find an even stronger positive correlation of 0.48 between C:N and C:P ratios. When we jointly analyze the three elemental contents, we find that N and P contents together explain 29% of the variance in C contents, that C and P together explain 31% of the variance in N contents, while P content is less well explained by C and N at 5%. These general relationships hide more complex patterns when data are analyzed at the taxonomic group level (Fig. 2, Table S3). Indeed three-way relationships are strongly variable among taxonomic groups (Fig. 2).

**Figure 2.**
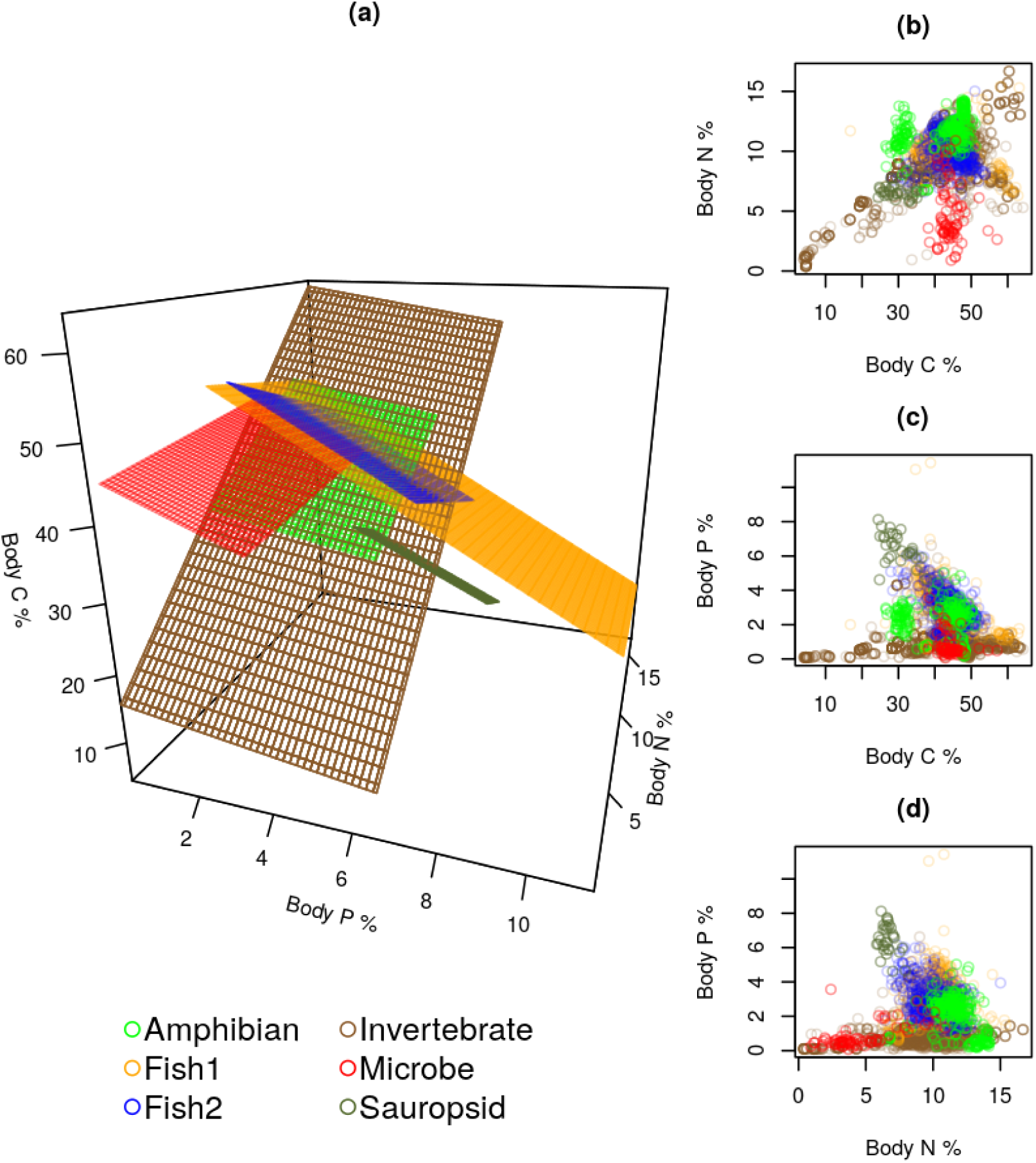
Co-variations in elemental content composition. (a) Stoichiometric spectrums computed as the best-fit planes of the linear regression of C content against N and P contents, for each taxonomical group separately. R^2^= 0.19 for amphibians, 0.38 for fishes-1 (Euteleosteomorpha), 0.39 for fishes-2 (Otomorpha), 0.47 for invertebrates, 0.13 for microbes, 0.29 for sauropsids. (b) Two-way relationships between C and N contents, raw data. (c) Two-way relationships between C and P contents, raw data. (d) Two-way relationships between N and P contents, raw data. There is no mammal data with simultaneous measurements of C, N and P contents.

### Stoichiometric drivers

Overall, the four stoichiometric drivers studied had significant effects on body stoichiometry. The full model including the four drivers assessed was always selected for the six stoichiometric traits (Table 1, Table S4). For four traits, C, N, C:P and N:P, it was the best model and was totaling almost all the Akaike weights. For P, it was the best model totaling 98 per cent of Akaike weights. And for C:N, it was the second best model, totaling 14 per cent of Akaike weights. For this last trait, a model without the habitat effect had a larger weight.

**Table 1.**
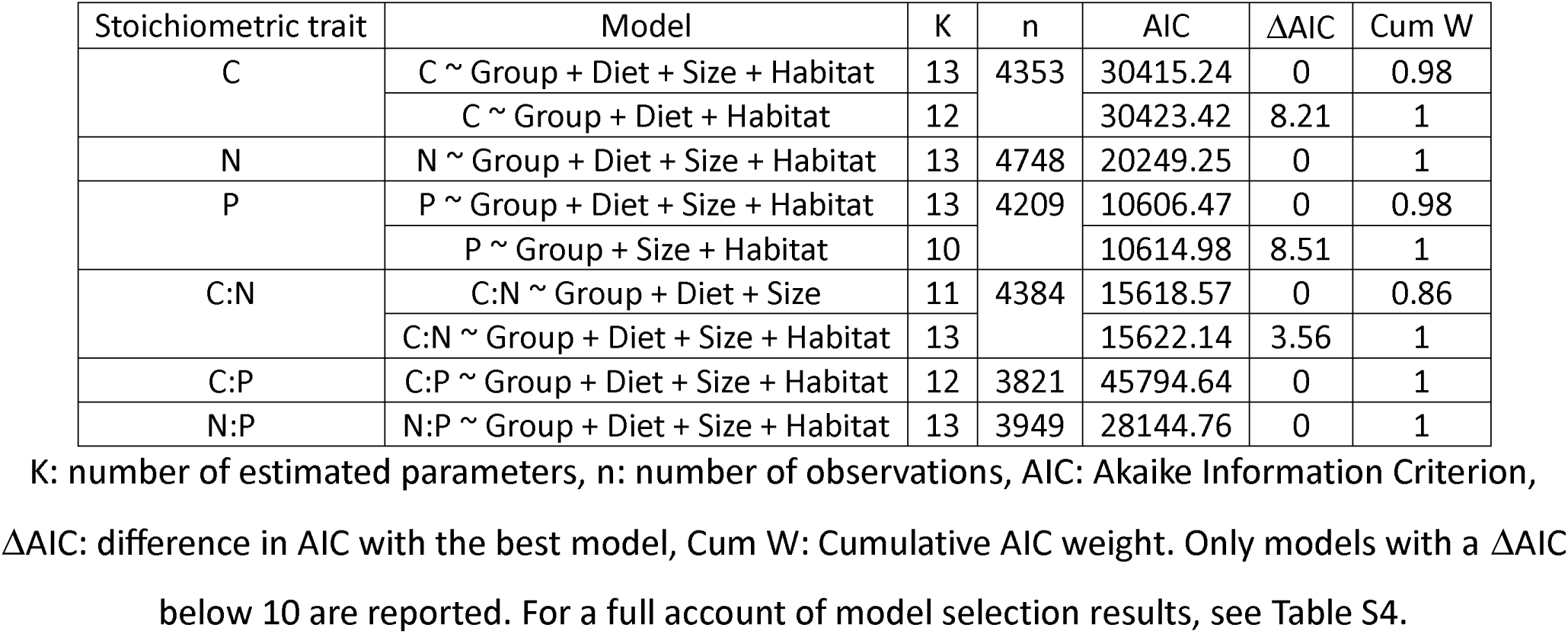
Stoichiometric drivers assessed by model selection.

Taxonomic group was the most influent driver among the four studied for all stoichiometric traits except carbon content (Fig. 3). This effect was especially large for P, C:P and N:P with partial eta-squared values above 0.1 (Fig. 3). Habitat was the strongest driver for C and the second most influent driver for four other stoichiometric traits (except C:N), while diet and body mass had only minor effects with partial eta-squared below 0.05 for all traits studied (Fig. 3). We also found that the effects of habitat, diet and body mass were group-specific, both in terms of magnitude and significance (Tables S5-10).

**Figure 3.**
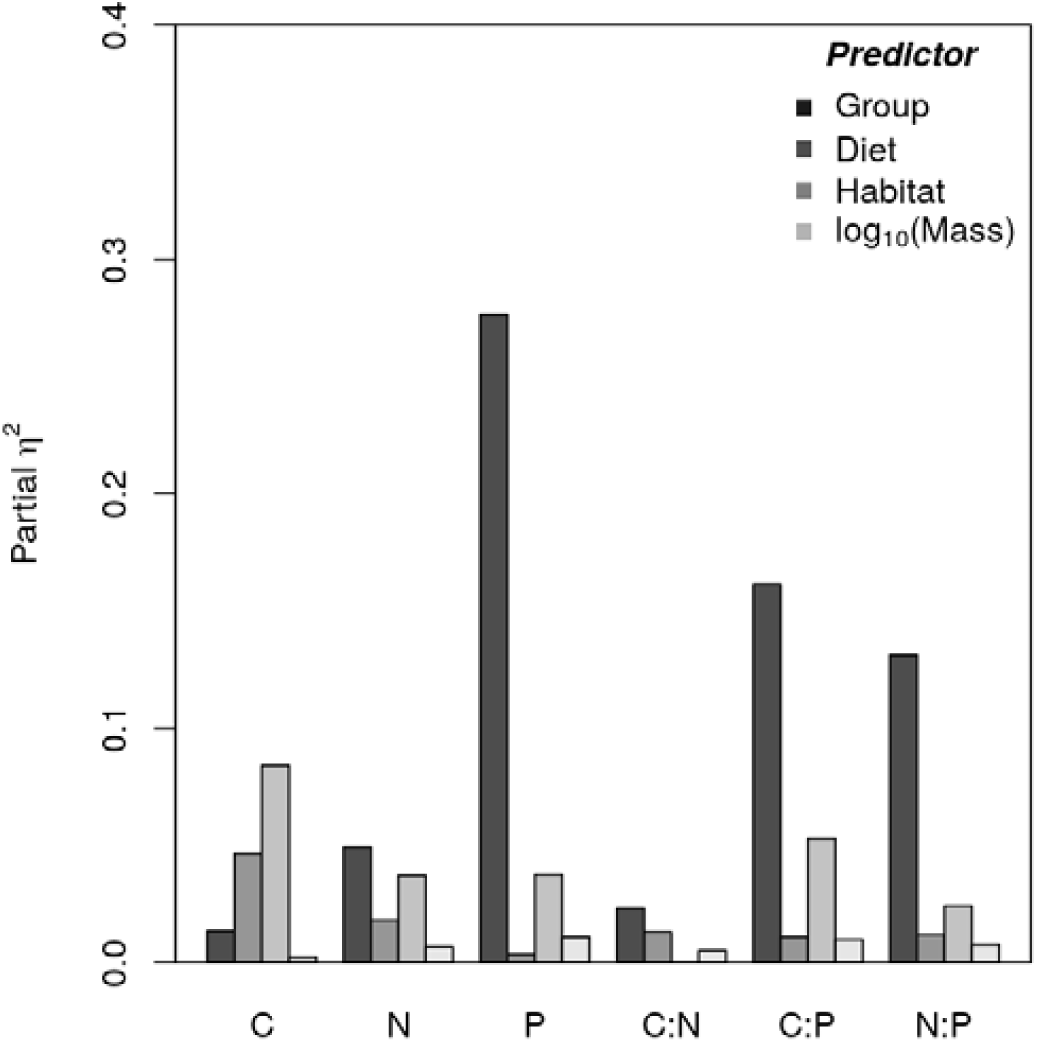
Magnitude of the stoichiometric drivers for the six stoichiometric traits assessed with partial eta-squared.

Our analysis based on sequential ANCOVAs confirmed these results (Fig. 4, Figs. S11-15). When looking at the results for P content (Fig. 4), we detected significant differences of sizable magnitude between the different taxonomic groups, while differences between diets, habitats and body mass were small once the effects of other drivers had been accounted for (Fig. 4). We obtained similar results for the other stoichiometric traits (Figs. S11-15).

**Figure 4.**
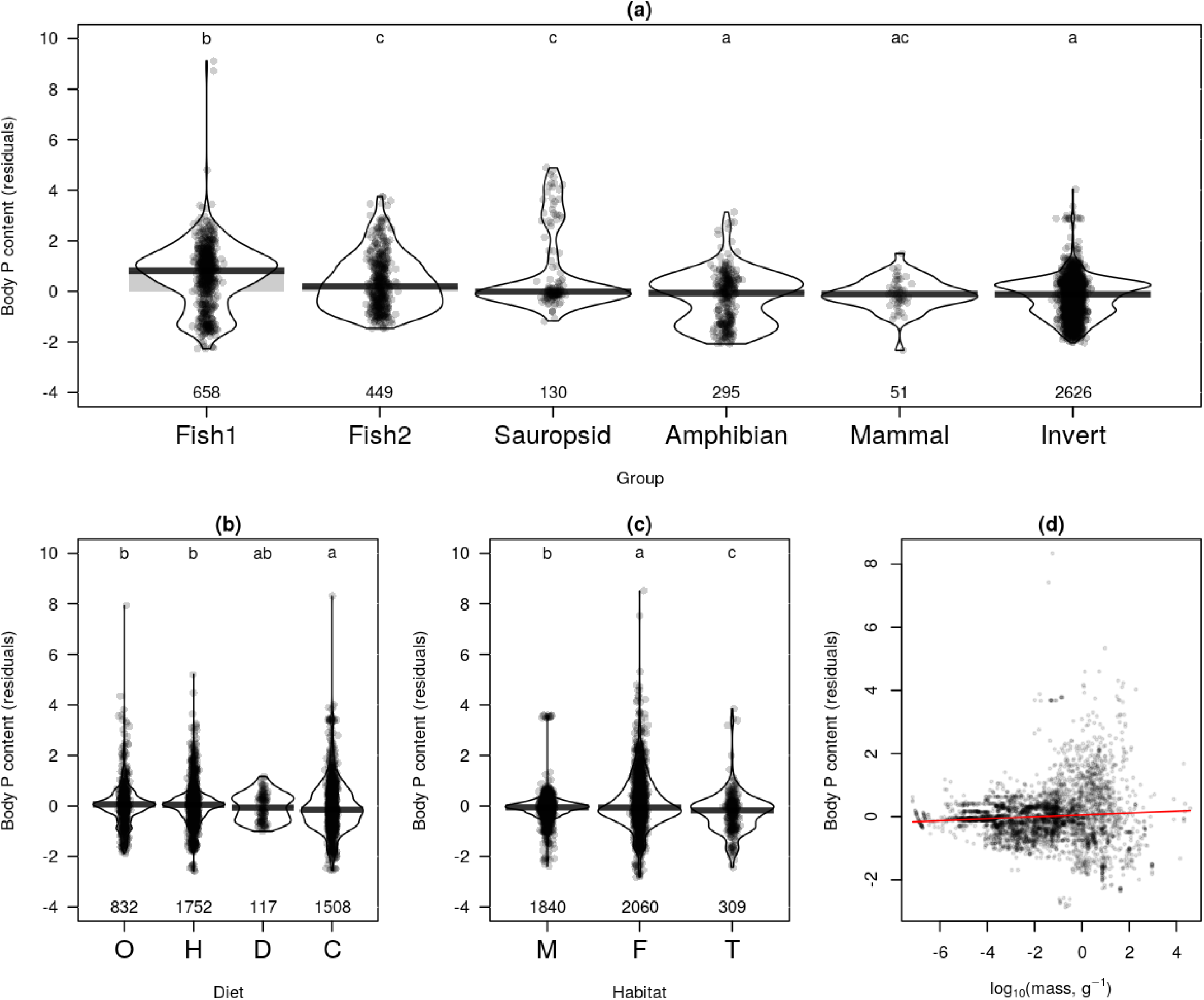
Magnitude of the stoichiometric drivers for P content assessed with sequential ANCOVAs. (a) taxonomic groups, (b) diet, (c) habitat and (d) body mass. Raw residuals (points) and distributions (kernel densities) are depicted with the number of observations (at the bottom for panels a-c). Different letters indicate significant differences with post-hoc Dunn’s test of multiple comparisons with Bonferroni corrections (panels a-c). The red line panel (d) show the prediction of body P content residuals regressed against log-transformed dry body masses (y = 0.03 + 0.05 x, R^2^ = 0.006, p < 0.001).

### Evolutionary history

We found a significant (p<0.05) phylogenetic signal with both methods and for the six stoichiometric traits (Table S16). Computed values of K were below 1 for the six stoichiometric traits, indicating that phylogenetically related species have less similar trait values than expected under a Brownian motion model of evolution (Fig. 5). Computed values of λ were close to 1, indicating that traits are as similar among species as awaited under a Brownian motion model of evolution (Fig. 5). Consequently, the two studied statistics provide somehow contradictory results, with K and λ suggesting a shallow and strong phylogenetic signal respectively.

**Figure 5.**
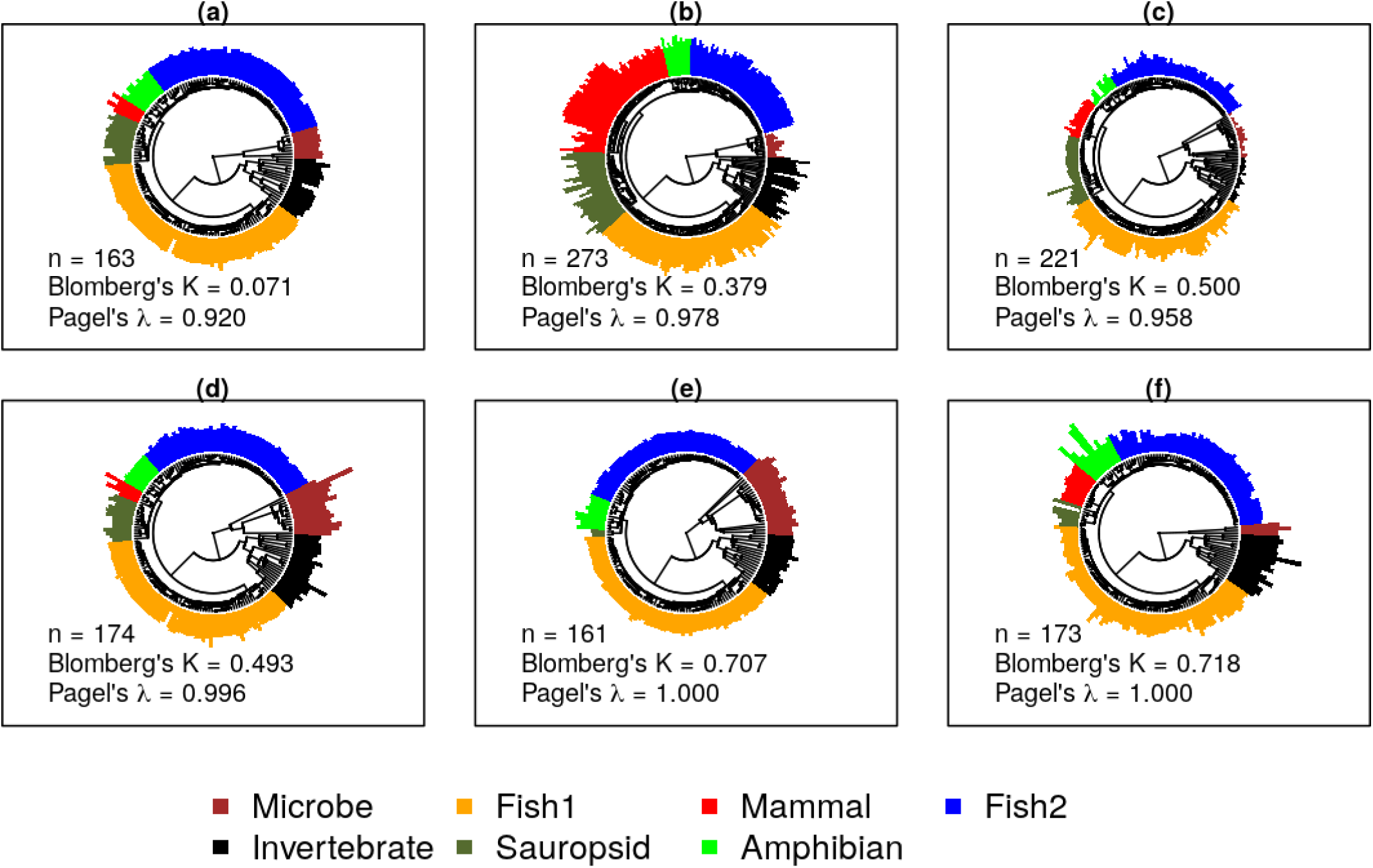
Phylogenetic variation of the six stoichiometric traits. (a) Body C content. (b) Body N content. (c) Body P content. (d) C:N ratio. (e) C:P ratio. (f) N:P ratio. Colored bars represent standardized values of the traits. Data were log-transformed for stoichiometric ratios. n: number of observations. See Tab. S16 for more details on phylogenetic signal analyses.

Our complementary analysis based on mixed models sheds light on this apparent contradiction by revealing the phylogenetic hierarchical structure of the variation of stoichiometric traits. Indeed, we found for the six stoichiometric traits large effects of taxonomic classifications at the three taxonomic levels considered (families, orders and classes, Table 2). For C, N and P contents, phylogenetic variation was strongly structured among classes, while variations among orders and families had lower and similar magnitudes (Table 2). For the C:N, C:P and N:P ratios, the largest variations were among families, followed by among-classes variations and among-orders variations (Table 2). Moreover, the predictive power of the linear mixed models of the six stoichiometric traits were largely greater when taxonomy was accounted for, as shown by the low marginal- and high conditional R^2^ values (Table 2).

**Table 2.** Magnitude of taxonomic effects on stoichiometric traits.

| Model | n | Marginal<br>R<sup>2</sup> | Conditional<br>R<sup>2</sup> | Random effects |  |  |  |
| --- | --- | --- | --- | --- | --- | --- | --- |
|  |  |  |  | Class | Order | Family | Residu |
| C ~ Diet + Habitat + Mass + Taxonomy (C/O/F) | 3746 | 0.008 | 0.948 | 0.866 | 0.013 | 0.070 | 0.05 |
| N ~ Diet + Habitat + Mass + Taxonomy (C/O/F) | 3999 | 0.014 | 0.909 | 0.668 | 0.120 | 0.119 | 0.09 |
| P ~ Diet + Habitat + Mass + Taxonomy (C/O/F) | 3462 | 0.081 | 0.723 | 0.421 | 0.101 | 0.176 | 0.30 |
| CN ~ Diet + Habitat + Mass + Taxonomy (C/O/F) | 3772 | 0.077 | 0.740 | 0.251 | 0.106 | 0.362 | 0.28 |
| CP ~ Diet + Habitat + Mass + Taxonomy (C/O/F) | 3242 | 0.007 | 0.796 | 0.266 | 0.168 | 0.360 | 0.20 |
| NP ~ Diet + Habitat + Mass + Taxonomy (C/O/F) | 3291 | 0.009 | 0.792 | 0.288 | 0.140 | 0.363 | 0.20 |

n: number of observations.

## DISCUSSION

### Analyzing interspecific variations among heterotrophs

To date, our compilation of body elemental contents of heterotrophs constitutes the most comprehensive resource to study the ecological stoichiometry of natural organisms with 17,848 independent observations on 1,491 species. Analyzing this database revealed widely variable carbon, nitrogen and phosphorus contents, as well as associated ratios (Fig. 1). We also evidenced sizeable levels of two and three-way co-variations of these elemental contents, but we found that these co-variations greatly vary between the different taxonomic groups considered (Fig. 2). The four assessed drivers significantly contribute to the observed interspecific variation in body elemental composition (Table 1), with taxonomic group being the strongest contributor, followed by habitat, diet and body mass (Fig. 3). Finally, we detected a significant variation of the six stoichiometric traits at the three nested taxonomic levels studied (classes, orders and families, Table 3). This leads to a significant but relatively modest phylogenetic signal (Fig. 5).

Consistently with available knowledge, vertebrates harbor larger levels of phosphorus content than invertebrates and microbes, since the skeleton contains a large amount of P (Hendrixson et al., 2007). Microbes present lower values of nitrogen content. Since our database contains many terrestrial fungi from the synthesis of Zhang & Elser (2017), this fact may principally highlight the necessary adaptation of such organisms to the frequent conditions of nitrogen limitation in the soil. This may also indicate the potential failure of the strict homeostasis assumption for these organisms (Danger et al. 2016). Mammals tend to have larger carbon and nitrogen contents. Large values of carbon content in mammals are probably due to their necessary lipid reserves to sustain their endothermy (Klaassen & Nolet, 2008). Finally, some invertebrate taxa present very low levels of carbon and nitrogen. These organisms belong to marine gelatinous taxa such as jellyfishes (Ikeda, 1974). Indeed, in these organisms, chloride, sodium and other minerals make up the bulk of organism dry mass (Khong et al., 2016). Besides, we observed a large between group variability in stoichiometric co-variations (Fig. 2). For instance, we found a strong positive co-variation between C and N content in invertebrates (Fig. 2a-b), while in fishes and sauropsids, the strongest covariation is negative between C and P contents (Fig. 2a,c). These sharp differences in stoichiometric co-variations among the taxonomic group considered suggest that these groups may face different physiological and life-history constraints. Indeed, insect exoskeletons are rich in chitin that contains both C and N, while vertebrate skeletons contain calcium phosphate and thus P.

### Predicting stoichiometric traits in heterotrophs

Our compiled database can be used to predict the main stoichiometric traits, i.e., body C, N and P contents and C:N, C:P and N:P ratios in heterotrophic organisms, accounting for their phylogenetic position, habitat, diet and body mass. Our analyses revealed that these four putative drivers significantly influence the stoichiometric traits studied (Fig.3). However, the magnitude of habitat, diet and body mass are small compared to the observed variations between taxonomic groups and to the remaining residual variations (Fig.4, Fig. S11-15). This modest magnitude of effects of the drivers studied may be partly explained by their relatively crude resolution in our database. Indeed, diet types may be more finely assessed thanks to discrete trophic levels, functional trophic guilds or isotopic niches (Majdi et al., 2018; McIntyre & Flecker, 2010). In the same vein, habitat definition may be more finely detailed so as to account for finer-scale habitat variation or environmental variables may be used instead (Zhang & Elser, 2017). Thus, we consider our analysis of stoichiometric drivers to be conservative, since it may underestimate the real effects of habitat and diet on interspecific variation in body elemental composition of heterotrophs. Complementing our current database with more accurate habitat and diet descriptions would thus constitute an interesting path for future progresses.

The larger effect of taxonomy on body stoichiometry (Fig. 3-4) with a significant phylogenetic signal for the six traits studied is also promising (Fig. 5). However, the large variability near the tips of the phylogeny (within families, Table 2) should make the imputation of stoichiometric traits from phylogenetic relatives a challenging task. We must recognize however that our phylogenetic analyses were based on relatively crude phylogenetic trees due to the taxonomic breadth of our database, so that more phylogenetically local analyses might provide clearer signals. Another related line of progress would consist in considering interactions between drivers in prediction models. The complete database was not amenable to such analyses, since it was not factorially complete. But analyses focused on particular phylogenetic groups would enable such more detailed treatments. Finally, on top of their effect on interspecific variation, habitat, diet and body mass are further likely to impact intraspecific variation in body elemental composition, as it has been recurrently evidenced in previous studies on specific taxonomic groups (Table S1, Lemmen et al., 2019). Our global database currently contains too few intraspecific repetitions to handle a meaningful global analysis on intraspecific variation in heterotrophs. This is clearly a line of progress that should be explored in the future.

## Acknowledgements

This research is funded by the French government IDEX-ISITE initiative 16-IDEX-0001 (CAP 20-25).

## DATA ACCESSIBILITY STATEMENT

- The compiled database will be deposited on Zenodo upon acceptance of publication.
- The R script used for the statistical analyses will be deposited on Github upon acceptance of publication.

