## Supplementary Information for "Body stoichiometry of heterotrophs: assessing drivers of interspecific variations in elemental composition"

**Table S1**: Major comprehensive studies about body elemental content in heterotrophs.

| Variables | Co-variates | Focused organisms | Main results | References |
| --- | --- | --- | --- | --- |
| C:N:P | Habitat (terrestrial vs freshwater) | zooplankton, insects | - C:N, C:P and N:P similar between terrestrial and freshwater herbivores | Elser et al. 2000 |
| N | Phylogeny, trophic level, mass, ontogeny | Insects | - Predators contain more N than herbivores, even when controlling for phylogeny and allometry - Phylogenetic variation in N content among herbivores at the ordinal group level. - Positive correlation between body mass and N content for predators, non-significant for herbivores. - No ontogenetic signal | Fagan et al. 2002 |
| C, N, P, C:N:P | Phylogeny | Fishes, amphibians | - Great variations in C:P and N:P among families, low variation in C, N and C:N - Within families, low variation in P content | Vanni et al. 2002 |
| P | Phylogeny, trophic level, mass | Insects, arachnids | - Inter- and intra-specific negative correlation between P content and mass - Predator and herbivore P content similar - Phylogenetic variation in P content at the ordinal group level. | Woods et al. 2004 |
| C, N, P, C:N:P | Phylogeny, habitat, mass, functional feeding group, season | Benthic insects, crustaceans and mollusks | - Strong phylogenetic variation in body elemental content and ratios at high taxonomic levels (classes and phyla) and at finer taxonomic levels. N and N:P highest for predators - Body mass explains little variation - Season explains little variation - Habitat explains little variation | Evans-White et al. 2005 |
| C, N, P | Mass, | Terrestrial insects (crickets) | - Negative correlation between body C and N or P, positive correlation between N and P. - Positive correlation between C and mass; negative correlation between mass and N or P. | Bertram et al. 2006 |
| C, N, P, C:N:P | Phylogeny, mass, habitat | Freshwater fishes | - High phylogenetic signal for P content, C:P and N:P and not for C, N and C:N - Negative correlation between C and P contents. - Intraspecific allometry is species-specific (sign and significance) - Across species, quadratic scaling for P and N:P but become non-significant when controlling for phylogeny - P content habitat-specific | Hendrixson et al. 2007 |
| N, P | Phylogeny, mass, trophic level | Detritivorous arthropods | - Phylogenetic variation for N and P at the ordinal group level. - No correlation with body mass - N content similar between detritivores and herbivores, higher in predators - P content independent of trophic level | Martinson et al 2008 |
| C, N, P, C:N:P | Taxonomy, trophic guild | Stream fishes | - Family and guild factors together explain 74% and 80% of inter-specific variations in N and P content, respectively - Guild only explain variation in N content, not P - Lower N content in primary consumers - Negative correlation between N and P contents | McIntyre and Flecker, 2010 |
| C, N, P, C:N:P | Phylogeny, trophic level, mass, ontogeny, habitat | Arthropods, lizards | - P content more variable than N content among and within species. - Significant phylogenetic signal for arthropods P content, but low variability explained (4%), N, C and C:N independent of phylogeny - Higher trophic levels have higher N, P, and lower C:N and N:P (arthropods) - Larger lizards contains more P and have lower N:P mass-scaling P content not significant when controlling for phylogeny in arthropods - Little habitat effect for most species. - Larger C:N in larvae than adults, no difference for N, P and C:P. | Gonzalez et al. 2011 |
| C, N, P, C:N:P | Mass, habitat | Crustaceans | - Species-dependent effect of habitat and mass on body stoichiometry | Arbaciauskas et al. 2013 |
| P | Phylogeny, trophic level, mass | Insects, spiders | - Predator have higher P contents than herbivores - Phylogeny explains variation in P content at the class, order, family and genus levels. | Wiesenborn 2013 |
| C, N, P, Ca, C:N:P | Mass, habitat | Salamanders (larvae) | - N, C:P and N:P decrease with mass - P and C:N increase with mass - Little effect of habitat | Milanovitch et al., 2015 |
| C, N, P, C:N:P | Phylum, functional guilds, environmental co-variates | Fungi | - C:P and N:P differ among phyla - C:N and N:P differ among guilds - Significant correlations of stoichiometric traits of Agaricomycete fungi with latitude, elevation, precipitation and temperature. | Zhang and Elser, 2017 |
| C, N, P, C:N:P | Phylogeny, taxonomy, trophic group, mass, habitat | Aquatic invertebrates (larvae) | - Taxonomy is a major predictor of body stoichiometry - Weak phylogenetic signal in elemental contents - Intra-specific variation in body stoichiometry is related to body mass with positive correlation with C:N, C:P and N:P and negative ones with N and P contents. - Controlling for phylogenetic, all interspecific allometric relationships disappear - Higher C,N, C:P and N:P and lower P and C:N in carnivores compared to detritivores - Habitat contributes to variation in body stoichiometry | Gonzalez et al., 2018 |

**Figure S2**: Description of the database. Panel (a) shows the total number of data entries and the number of species per group. Panel (b) shows the fraction of diet type per group. Panel (c) shows the total number of data entries per habitat and the number of different species per habitat. Panel (d) shows the density distribution of the mass variable per group. Invert: Invertebrate, Fish1: Euteleosteomorpha, Fish2: Otomorpha. Note the log axes on panels (a) and (d).
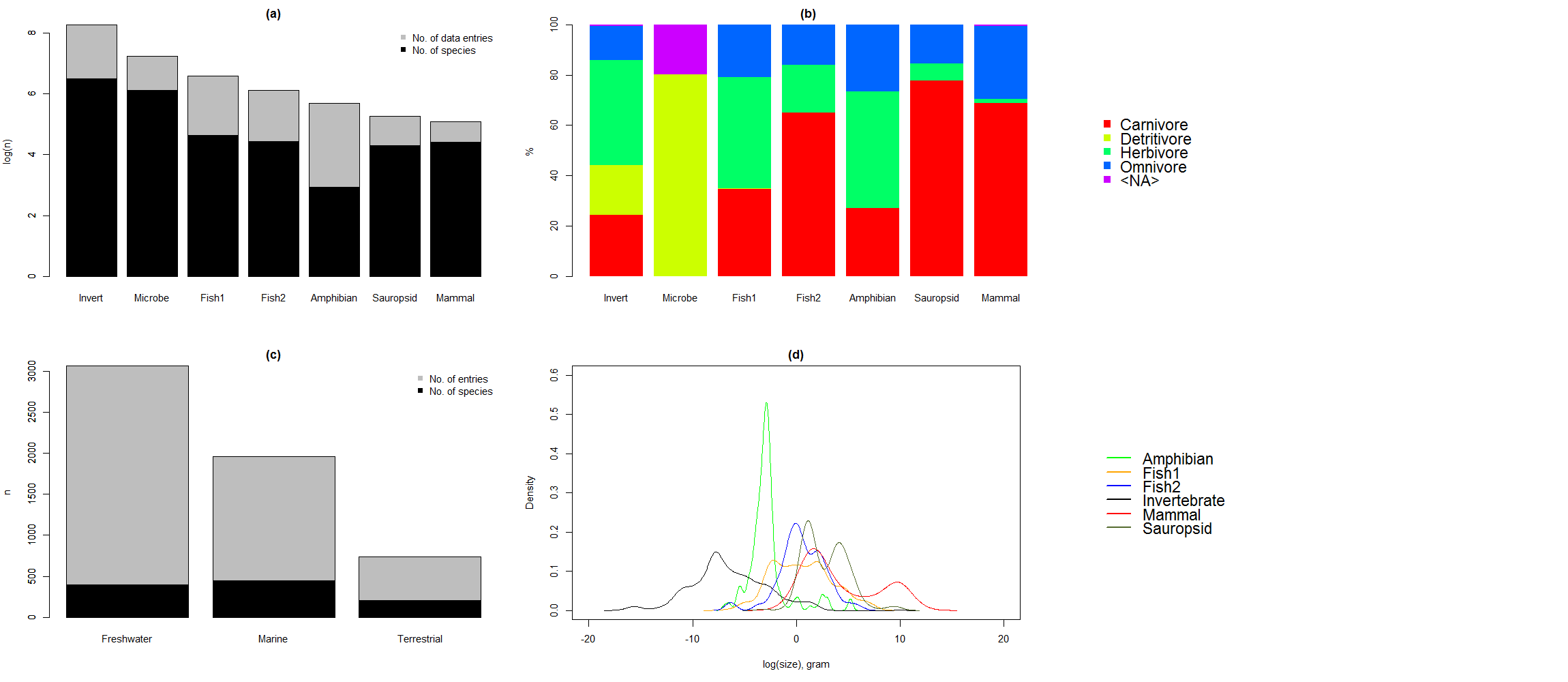

**Table S3**: Pearson’s product moment correlation coefficient between body C and N contents (a), body C and P contents (b), body N and P contents (c), for each group of heterotrophs. P-values < 0.05 are shown in bold. n: number of observations.

|  | (a) C-N | | | (b) C-P | | | (c) N-P | | |
| --- | --- | --- | --- | --- | --- | --- | --- | --- | --- |
| Group | n | Pearson's r | p-value | n | Pearson's r | p-value | n | Pearson's r | p-value |
| Amphibian | 258 | 0.41 | **< 0.001** | 258 | -0.27 | **< 0.001** | 293 | -0.19 | **0.001** |
| Fish1 - Euteleosteomorpha | 616 | -0.2 | **< 0.001** | 606 | -0.6 | **< 0.001** | 675 | 0.21 | **< 0.001** |
| Fish2 - Otomorpha | 452 | -0.11 | **0.023** | 452 | -0.53 | **< 0.001** | 456 | -0.37 | **< 0.001** |
| Invertebrate | 3399 | 0.64 | **< 0.001** | 2478 | 0.08 | **< 0.001** | 2540 | 0.18 | **< 0.001** |
| Mammal | 10 | -0.9 | **< 0.001** | - | - | - | 31 | 0.02 | 0.927 |
| Microbe | 341 | -0.01 | 0.904 | 89 | -0.25 | **0.017** | 250 | 0.5 | **< 0.001** |
| Sauropsid | 59 | 0.74 | **< 0.001** | 31 | -0.5 | **0.004** | 45 | -0.8 | **< 0.001** |

**Table S4**: Full account of model selection results.

| Stoichiometric trait | Model | K | AIC | Delta_AIC | ModelLik | AICWt | LL | Cum.Wt |
| --- | --- | --- | --- | --- | --- | --- | --- | --- |
| C | C ~ Group + Diet + Size + Habitat | 13 | 30415.24 | 0.00 | 1.00 | 0.98 | -15194.62 | 0.98 |
|  | C ~ Group + Diet + Habitat | 12 | 30423.45 | 8.21 | 0.02 | 0.02 | -15199.72 | 1.00 |
|  | C ~ Diet + Size + Habitat | 8 | 30464.58 | 49.35 | 0.00 | 0.00 | -15224.29 | 1.00 |
|  | C ~ Diet + Habitat | 7 | 30489.12 | 73.89 | 0.00 | 0.00 | -15237.56 | 1.00 |
|  | C ~ Group + Size + Habitat | 10 | 30615.98 | 200.74 | 0.00 | 0.00 | -15297.99 | 1.00 |
|  | C ~ Group + Habitat | 9 | 30630.47 | 215.23 | 0.00 | 0.00 | -15306.23 | 1.00 |
|  | C ~ Size + Habitat | 5 | 30719.90 | 304.67 | 0.00 | 0.00 | -15354.95 | 1.00 |
|  | C ~ Group + Diet | 10 | 30792.63 | 377.40 | 0.00 | 0.00 | -15386.32 | 1.00 |
|  | C ~ Group + Diet + Size | 11 | 30794.18 | 378.94 | 0.00 | 0.00 | -15386.09 | 1.00 |
|  | C ~ Habitat | 4 | 30804.79 | 389.55 | 0.00 | 0.00 | -15398.40 | 1.00 |
|  | C ~ Diet + Size | 6 | 30854.82 | 439.58 | 0.00 | 0.00 | -15421.41 | 1.00 |
|  | C ~ Diet | 5 | 30880.24 | 465.00 | 0.00 | 0.00 | -15435.12 | 1.00 |
|  | C ~ Group | 7 | 31128.41 | 713.17 | 0.00 | 0.00 | -15557.20 | 1.00 |
|  | C ~ Group + Size | 8 | 31129.94 | 714.71 | 0.00 | 0.00 | -15556.97 | 1.00 |
|  | C ~ Size | 3 | 31173.67 | 758.43 | 0.00 | 0.00 | -15583.84 | 1.00 |
| N | N ~ Group + Diet + Size + Habitat | 13 | 20249.25 | 0.00 | 1.00 | 1.00 | -10111.63 | 1.00 |
|  | N ~ Group + Diet + Habitat | 12 | 20279.04 | 29.78 | 0.00 | 0.00 | -10127.52 | 1.00 |
|  | N ~ Group + Size + Habitat | 10 | 20329.46 | 80.21 | 0.00 | 0.00 | -10154.73 | 1.00 |
|  | N ~ Group + Habitat | 9 | 20359.86 | 110.60 | 0.00 | 0.00 | -10170.93 | 1.00 |
|  | N ~ Group + Diet + Size | 11 | 20425.63 | 176.38 | 0.00 | 0.00 | -10201.82 | 1.00 |
|  | N ~ Group + Diet | 10 | 20441.57 | 192.32 | 0.00 | 0.00 | -10210.78 | 1.00 |
|  | N ~ Diet + Size + Habitat | 8 | 20479.94 | 230.69 | 0.00 | 0.00 | -10231.97 | 1.00 |
|  | N ~ Diet + Habitat | 7 | 20491.89 | 242.64 | 0.00 | 0.00 | -10238.95 | 1.00 |
|  | N ~ Size + Habitat | 5 | 20530.46 | 281.21 | 0.00 | 0.00 | -10260.23 | 1.00 |
|  | N ~ Group + Size | 8 | 20539.13 | 289.88 | 0.00 | 0.00 | -10261.57 | 1.00 |
|  | N ~ Habitat | 4 | 20543.24 | 293.99 | 0.00 | 0.00 | -10267.62 | 1.00 |
|  | N ~ Group | 7 | 20558.34 | 309.09 | 0.00 | 0.00 | -10272.17 | 1.00 |
|  | N ~ Diet + Size | 6 | 20844.86 | 595.61 | 0.00 | 0.00 | -10416.43 | 1.00 |
|  | N ~ Diet | 5 | 20880.95 | 631.70 | 0.00 | 0.00 | -10435.48 | 1.00 |
|  | N ~ Size | 3 | 20887.12 | 637.87 | 0.00 | 0.00 | -10440.56 | 1.00 |
| P | P ~ Group + Diet + Size + Habitat | 13 | 10606.47 | 0.00 | 1.00 | 0.99 | -5290.24 | 0.99 |
|  | P ~ Group + Size + Habitat | 10 | 10614.98 | 8.51 | 0.01 | 0.01 | -5297.49 | 1.00 |
|  | P ~ Group + Diet + Habitat | 12 | 10649.76 | 43.28 | 0.00 | 0.00 | -5312.88 | 1.00 |
|  | P ~ Group + Habitat | 9 | 10653.61 | 47.13 | 0.00 | 0.00 | -5317.80 | 1.00 |
|  | P ~ Group + Diet + Size | 11 | 10764.29 | 157.82 | 0.00 | 0.00 | -5371.15 | 1.00 |
|  | P ~ Group + Size | 8 | 10783.78 | 177.31 | 0.00 | 0.00 | -5383.89 | 1.00 |
|  | P ~ Group + Diet | 10 | 10841.10 | 234.63 | 0.00 | 0.00 | -5410.55 | 1.00 |
|  | P ~ Group | 7 | 10854.63 | 248.15 | 0.00 | 0.00 | -5420.31 | 1.00 |
|  | P ~ Diet + Size + Habitat | 8 | 11958.99 | 1352.51 | 0.00 | 0.00 | -5971.49 | 1.00 |
|  | P ~ Size + Habitat | 5 | 12088.76 | 1482.28 | 0.00 | 0.00 | -6039.38 | 1.00 |
|  | P ~ Diet + Size | 6 | 12509.51 | 1903.04 | 0.00 | 0.00 | -6248.76 | 1.00 |
|  | P ~ Diet + Habitat | 7 | 12583.64 | 1977.16 | 0.00 | 0.00 | -6284.82 | 1.00 |
|  | P ~ Size | 3 | 12595.24 | 1988.76 | 0.00 | 0.00 | -6294.62 | 1.00 |
|  | P ~ Habitat | 4 | 12902.08 | 2295.61 | 0.00 | 0.00 | -6447.04 | 1.00 |
|  | P ~ Diet | 5 | 13891.80 | 3285.33 | 0.00 | 0.00 | -6940.90 | 1.00 |
| CN | CN ~ Group + Diet + Size | 11 | 15618.58 | 0.00 | 1.00 | 0.86 | -7798.29 | 0.86 |
|  | CN ~ Group + Diet + Size + Habitat | 13 | 15622.14 | 3.56 | 0.17 | 0.14 | -7798.07 | 1.00 |
|  | CN ~ Group + Diet | 10 | 15639.22 | 20.64 | 0.00 | 0.00 | -7809.61 | 1.00 |
|  | CN ~ Group + Diet + Habitat | 12 | 15642.58 | 24.01 | 0.00 | 0.00 | -7809.29 | 1.00 |
|  | CN ~ Group + Size + Habitat | 10 | 15673.72 | 55.15 | 0.00 | 0.00 | -7826.86 | 1.00 |
|  | CN ~ Group + Size | 8 | 15676.28 | 57.70 | 0.00 | 0.00 | -7830.14 | 1.00 |
|  | CN ~ Group | 7 | 15701.22 | 82.64 | 0.00 | 0.00 | -7843.61 | 1.00 |
|  | CN ~ Group + Habitat | 9 | 15702.04 | 83.46 | 0.00 | 0.00 | -7842.02 | 1.00 |
|  | CN ~ Diet + Size + Habitat | 8 | 15714.96 | 96.39 | 0.00 | 0.00 | -7849.48 | 1.00 |
|  | CN ~ Diet + Size | 6 | 15728.36 | 109.78 | 0.00 | 0.00 | -7858.18 | 1.00 |
|  | CN ~ Diet + Habitat | 7 | 15731.07 | 112.49 | 0.00 | 0.00 | -7858.53 | 1.00 |
|  | CN ~ Diet | 5 | 15733.23 | 114.66 | 0.00 | 0.00 | -7861.62 | 1.00 |
|  | CN ~ Size + Habitat | 5 | 15791.79 | 173.22 | 0.00 | 0.00 | -7890.90 | 1.00 |
|  | CN ~ Habitat | 4 | 15804.68 | 186.10 | 0.00 | 0.00 | -7898.34 | 1.00 |
|  | CN ~ Size | 3 | 15825.37 | 206.79 | 0.00 | 0.00 | -7909.68 | 1.00 |
| CP | CP ~ Group + Diet + Size + Habitat | 12 | 45794.64 | 0.00 | 1.00 | 1.00 | -22885.32 | 1.00 |
|  | CP ~ Group + Diet + Habitat | 11 | 45829.99 | 35.35 | 0.00 | 0.00 | -22904.00 | 1.00 |
|  | CP ~ Group + Size + Habitat | 9 | 45830.02 | 35.39 | 0.00 | 0.00 | -22906.01 | 1.00 |
|  | CP ~ Group + Habitat | 8 | 45856.41 | 61.77 | 0.00 | 0.00 | -22920.20 | 1.00 |
|  | CP ~ Group + Diet + Size | 10 | 45998.63 | 204.00 | 0.00 | 0.00 | -22989.32 | 1.00 |
|  | CP ~ Group + Diet | 9 | 46003.70 | 209.06 | 0.00 | 0.00 | -22992.85 | 1.00 |
|  | CP ~ Group + Size | 7 | 46028.79 | 234.15 | 0.00 | 0.00 | -23007.39 | 1.00 |
|  | CP ~ Group | 6 | 46030.14 | 235.50 | 0.00 | 0.00 | -23009.07 | 1.00 |
|  | CP ~ Diet + Size + Habitat | 8 | 46458.61 | 663.97 | 0.00 | 0.00 | -23221.30 | 1.00 |
|  | CP ~ Diet + Size | 6 | 46459.18 | 664.54 | 0.00 | 0.00 | -23223.59 | 1.00 |
|  | CP ~ Size + Habitat | 5 | 46486.59 | 691.95 | 0.00 | 0.00 | -23238.29 | 1.00 |
|  | CP ~ Size | 3 | 46491.26 | 696.62 | 0.00 | 0.00 | -23242.63 | 1.00 |
|  | CP ~ Diet + Habitat | 7 | 46773.45 | 978.81 | 0.00 | 0.00 | -23379.73 | 1.00 |
|  | CP ~ Habitat | 4 | 46822.11 | 1027.47 | 0.00 | 0.00 | -23407.06 | 1.00 |
|  | CP ~ Diet | 5 | 46848.07 | 1053.43 | 0.00 | 0.00 | -23419.04 | 1.00 |
| NP | NP ~ Group + Diet + Size + Habitat | 13 | 28144.76 | 0.00 | 1.00 | 1.00 | -14059.38 | 1.00 |
|  | NP ~ Group + Diet + Habitat | 12 | 28173.30 | 28.54 | 0.00 | 0.00 | -14074.65 | 1.00 |
|  | NP ~ Group + Size + Habitat | 10 | 28185.12 | 40.36 | 0.00 | 0.00 | -14082.56 | 1.00 |
|  | NP ~ Group + Habitat | 9 | 28205.62 | 60.86 | 0.00 | 0.00 | -14093.81 | 1.00 |
|  | NP ~ Group + Diet + Size | 11 | 28238.01 | 93.25 | 0.00 | 0.00 | -14108.00 | 1.00 |
|  | NP ~ Group + Diet | 10 | 28247.98 | 103.22 | 0.00 | 0.00 | -14113.99 | 1.00 |
|  | NP ~ Group + Size | 8 | 28270.52 | 125.76 | 0.00 | 0.00 | -14127.26 | 1.00 |
|  | NP ~ Group | 7 | 28275.97 | 131.21 | 0.00 | 0.00 | -14130.98 | 1.00 |
|  | NP ~ Diet + Size | 6 | 28686.73 | 541.97 | 0.00 | 0.00 | -14337.37 | 1.00 |
|  | NP ~ Diet + Size + Habitat | 8 | 28690.70 | 545.94 | 0.00 | 0.00 | -14337.35 | 1.00 |
|  | NP ~ Size | 3 | 28704.90 | 560.14 | 0.00 | 0.00 | -14349.45 | 1.00 |
|  | NP ~ Size + Habitat | 5 | 28708.33 | 563.57 | 0.00 | 0.00 | -14349.17 | 1.00 |
|  | NP ~ Diet + Habitat | 7 | 29009.96 | 865.20 | 0.00 | 0.00 | -14497.98 | 1.00 |
|  | NP ~ Habitat | 4 | 29047.74 | 902.98 | 0.00 | 0.00 | -14519.87 | 1.00 |
|  | NP ~ Diet | 5 | 29116.32 | 971.56 | 0.00 | 0.00 | -14553.16 | 1.00 |

**Table S5**: Model parameters for body C content as a function of diet, habitat and body size for each group of heterotrophs. *Fish1*: Euteleosteomorpha, *Fish2*: Otomorpha, *DietC*: carnivore, *DietD*: detritivore, *DietH*: herbivore, *DietO*: omnivore, *HabitatFreshwater*: freshwater, *HabitatMarine*: marine, *HabitatTerrestrial*: terrestrial, *Size*: log_10_(dry mass). P-values < 0.05 are shown in bold.

| Group | Level of comparison | Parameter | Estimate | Std. Error | t value | Pr(>\|t\|) |
| --- | --- | --- | --- | --- | --- | --- |
| Amphibian | DietC | (Intercept) | 41.268 | 0.742 | 55.625 | **< 0.001** |
|  |  | DietH | 5.953 | 1.102 | 5.404 | **< 0.001** |
|  |  | DietO | 8.211 | 1.900 | 4.321 | **< 0.001** |
|  |  | Size | 0.742 | 0.539 | 1.375 | 0.170 |
| Fish1 | DietC HabitatFreshwater | (Intercept) | 42.809 | 0.263 | 162.504 | **< 0.001** |
|  |  | DietH | -3.413 | 0.809 | -4.219 | **< 0.001** |
|  |  | DietO | -1.258 | 0.604 | -2.083 | **0.038** |
|  |  | HabitatMarine | 5.220 | 0.823 | 6.343 | **< 0.001** |
|  |  | Size | 0.933 | 0.208 | 4.492 | **< 0.001** |
| Fish2 | DietC HabitatFreshwater | (Intercept) | 45.011 | 0.347 | 129.566 | **< 0.001** |
|  |  | DietH | -1.740 | 0.461 | -3.773 | **< 0.001** |
|  |  | DietO | -0.966 | 0.588 | -1.643 | 0.101 |
|  |  | HabitatMarine | 1.449 | 2.150 | 0.674 | 0.501 |
|  |  | Size | 0.592 | 0.221 | 2.681 | **0.008** |
| Invertebrate | DietC HabitatFreshwater | (Intercept) | 41.742 | 0.440 | 94.921 | **< 0.001** |
|  |  | DietD | 3.833 | 0.600 | 6.384 | **< 0.001** |
|  |  | DietH | 6.602 | 0.390 | 16.907 | **< 0.001** |
|  |  | DietO | 3.718 | 0.503 | 7.387 | **< 0.001** |
|  |  | HabitatMarine | -8.317 | 0.381 | -21.845 | **< 0.001** |
|  |  | HabitatTerrestrial | -1.488 | 1.113 | -1.336 | 0.182 |
|  |  | Size | -0.619 | 0.103 | -5.993 | **< 0.001** |
| Mammal | DietC HabitatMarine | (Intercept) | 20.203 | 17.462 | 1.157 | 0.300 |
|  |  | DietH | 6.622 | 6.314 | 1.049 | 0.342 |
|  |  | DietO | 0.793 | 8.334 | 0.095 | 0.928 |
|  |  | HabitatTerrestrial | -14.438 | 8.296 | -1.740 | 0.142 |
|  |  | Size | 9.540 | 3.754 | 2.541 | 0.052 |
| Sauropsid | DietC HabitatFreshwater | (Intercept) | 22.082 | 2.110 | 10.464 | **< 0.001** |
|  |  | DietH | -3.792 | 3.562 | -1.065 | 0.292 |
|  |  | DietO | -0.317 | 1.210 | -0.262 | 0.794 |
|  |  | HabitatMarine | 5.401 | 5.132 | 1.052 | 0.297 |
|  |  | HabitatTerrestrial | 12.933 | 1.264 | 10.229 | **< 0.001** |
|  |  | Size | 4.889 | 1.063 | 4.598 | **< 0.001** |

**Table S6**: Model parameters for body N content as a function of diet, habitat and body size for each group of heterotrophs. *Fish1*: Euteleosteomorpha, *Fish2*: Otomorpha, *DietC*: carnivore, *DietD*: detritivore, *DietH*: herbivore, *DietO*: omnivore, *HabitatFreshwater*: freshwater, *HabitatMarine*: marine, *HabitatTerrestrial*: terrestrial, *Size*: log_10_(dry mass). P-values < 0.05 are shown in bold.

| Group | Level of comparison | Parameter | Estimate | Std. Error | t value | Pr(>\|t\|) |
| --- | --- | --- | --- | --- | --- | --- |
| Amphibian | DietC HabitatFreshwater | (Intercept) | 11.347 | 0.193 | 58.761 | **< 0.001** |
|  |  | DietH | 0.884 | 0.288 | 3.073 | **0.002** |
|  |  | DietO | 0.472 | 0.304 | 1.553 | 0.122 |
|  |  | HabitatTerrestrial | -0.046 | 0.697 | -0.065 | 0.948 |
|  |  | Size | -0.073 | 0.139 | -0.521 | 0.603 |
| Fish1 | DietC HabitatFreshwater | (Intercept) | 10.498 | 0.060 | 173.927 | **< 0.001** |
|  |  | DietH | -0.663 | 0.191 | -3.468 | **0.001** |
|  |  | DietO | -0.639 | 0.135 | -4.724 | **< 0.001** |
|  |  | HabitatMarine | -1.378 | 0.136 | -10.160 | **< 0.001** |
|  |  | Size | 0.116 | 0.040 | 2.892 | **0.004** |
| Fish2 | DietC HabitatFreshwater | (Intercept) | 10.136 | 0.108 | 94.036 | **< 0.001** |
|  |  | DietH | -0.411 | 0.143 | -2.871 | **0.004** |
|  |  | DietO | -0.046 | 0.182 | -0.252 | 0.801 |
|  |  | HabitatMarine | 1.650 | 0.667 | 2.473 | **0.014** |
|  |  | Size | 0.189 | 0.068 | 2.762 | **0.006** |
| Invertebrate | DietC HabitatFreshwater | (Intercept) | 9.107 | 0.112 | 81.498 | **< 0.001** |
|  |  | DietD | 0.503 | 0.150 | 3.362 | **0.001** |
|  |  | DietH | 1.094 | 0.100 | 10.958 | **< 0.001** |
|  |  | DietO | 0.601 | 0.130 | 4.628 | **< 0.001** |
|  |  | HabitatMarine | -0.981 | 0.097 | -10.145 | **< 0.001** |
|  |  | HabitatTerrestrial | 0.244 | 0.288 | 0.847 | 0.397 |
|  |  | Size | -0.148 | 0.026 | -5.603 | **< 0.001** |
| Mammal | DietC HabitatMarine | (Intercept) | 10.606 | 0.871 | 12.179 | **< 0.001** |
|  |  | DietH | -1.265 | 0.606 | -2.090 | **0.038** |
|  |  | DietO | -3.796 | 0.640 | -5.934 | **< 0.001** |
|  |  | HabitatTerrestrial | 4.281 | 0.927 | 4.620 | **< 0.001** |
|  |  | Size | -0.634 | 0.161 | -3.936 | **< 0.001** |
| Sauropsid | DietC HabitatFreshwater | (Intercept) | 7.426 | 0.636 | 11.672 | **< 0.001** |
|  |  | DietH | 3.499 | 1.567 | 2.233 | **0.028** |
|  |  | DietO | -0.614 | 0.444 | -1.383 | 0.170 |
|  |  | HabitatMarine | 5.477 | 2.204 | 2.485 | **0.015** |
|  |  | HabitatTerrestrial | 2.815 | 0.440 | 6.401 | **< 0.001** |
|  |  | Size | 0.370 | 0.265 | 1.397 | 0.166 |

**Table S7**: Model parameters for body P content as a function of diet, habitat and body size for each group of heterotrophs. *Fish1*: Euteleosteomorpha, *Fish2*: Otomorpha, *DietC*: carnivore, *DietD*: detritivore, *DietH*: herbivore, *DietO*: omnivore, *HabitatFreshwater*: freshwater, *HabitatMarine*: marine, *HabitatTerrestrial*: terrestrial, *Size*: log_10_(dry mass). P-values < 0.05 are shown in bold.

| Group | Level of comparison | Parameter | Estimate | Std. Error | t value | Pr(>\|t\|) |
| --- | --- | --- | --- | --- | --- | --- |
| Amphibian | DietC HabitatFreshwater | (Intercept) | 2.581 | 0.071 | 36.432 | **< 0.001** |
|  |  | DietH | -1.962 | 0.105 | -18.603 | **< 0.001** |
|  |  | DietO | -1.584 | 0.110 | -14.404 | **< 0.001** |
|  |  | HabitatTerrestrial | 1.995 | 0.242 | 8.250 | **< 0.001** |
|  |  | Size | 0.007 | 0.051 | 0.142 | 0.887 |
| Fish1 | DietC HabitatFreshwater | (Intercept) | 3.259 | 0.054 | 60.478 | **< 0.001** |
|  |  | DietH | 0.119 | 0.169 | 0.705 | 0.481 |
|  |  | DietO | 0.332 | 0.120 | 2.772 | **0.006** |
|  |  | HabitatMarine | -1.186 | 0.144 | -8.250 | **< 0.001** |
|  |  | Size | -0.227 | 0.039 | -5.771 | **< 0.001** |
| Fish2 | DietC HabitatFreshwater | (Intercept) | 2.932 | 0.082 | 35.555 | **< 0.001** |
|  |  | DietH | 0.039 | 0.109 | 0.361 | 0.718 |
|  |  | DietO | -0.162 | 0.139 | -1.161 | 0.246 |
|  |  | HabitatMarine | -0.820 | 0.510 | -1.607 | 0.109 |
|  |  | Size | -0.073 | 0.052 | -1.396 | 0.163 |
| Invertebrate | DietC HabitatFreshwater | (Intercept) | 1.029 | 0.032 | 32.071 | **< 0.001** |
|  |  | DietD | 0.059 | 0.061 | 0.969 | 0.333 |
|  |  | DietH | 0.338 | 0.027 | 12.317 | **< 0.001** |
|  |  | DietO | 0.303 | 0.034 | 8.916 | **< 0.001** |
|  |  | HabitatMarine | -0.110 | 0.026 | -4.176 | **< 0.001** |
|  |  | HabitatTerrestrial | -0.041 | 0.052 | -0.783 | 0.434 |
|  |  | Size | 0.106 | 0.007 | 15.617 | **< 0.001** |
| Mammal | DietC | (Intercept) | 1.931 | 0.272 | 7.109 | **< 0.001** |
|  |  | DietH | 0.304 | 0.274 | 1.108 | 0.273 |
|  |  | DietO | -0.014 | 0.272 | -0.053 | 0.958 |
|  |  | Size | 0.050 | 0.052 | 0.968 | 0.338 |
| Sauropsid | DietC HabitatFreshwater | (Intercept) | 6.178 | 0.246 | 25.119 | **< 0.001** |
|  |  | DietO | -0.071 | 0.141 | -0.506 | 0.614 |
|  |  | HabitatTerrestrial | -4.143 | 0.196 | -21.121 | **< 0.001** |
|  |  | Size | 0.065 | 0.122 | 0.530 | 0.597 |

**Table S8**: Model parameters for body C:N ratio as a function of diet, habitat and body size for each group of heterotrophs. *Fish1*: Euteleosteomorpha, *Fish2*: Otomorpha, *DietC*: carnivore, *DietD*: detritivore, *DietH*: herbivore, *DietO*: omnivore, *HabitatFreshwater*: freshwater, *HabitatMarine*: marine, *HabitatTerrestrial*: terrestrial, *Size*: log_10_(dry mass). P-values < 0.05 are shown in bold.

| Group | Level of comparison | Parameter | Estimate | Std. Error | t value | Pr(>\|t\|) |
| --- | --- | --- | --- | --- | --- | --- |
| Amphibian | DietC | (Intercept) | 4.252 | 0.082 | 51.870 | **< 0.001** |
|  |  | DietH | 0.380 | 0.122 | 3.119 | **0.002** |
|  |  | DietO | 0.094 | 0.210 | 0.449 | 0.654 |
|  |  | Size | 0.104 | 0.060 | 1.753 | 0.081 |
| Fish1 | DietC HabitatFreshwater | (Intercept) | 4.809 | 0.057 | 84.449 | **< 0.001** |
|  |  | DietH | -0.069 | 0.175 | -0.397 | 0.691 |
|  |  | DietO | -0.018 | 0.131 | -0.138 | 0.891 |
|  |  | HabitatMarine | 1.572 | 0.178 | 8.839 | **< 0.001** |
|  |  | Size | -0.031 | 0.045 | -0.701 | 0.483 |
| Fish2 | DietC HabitatFreshwater | (Intercept) | 5.266 | 0.076 | 69.248 | **< 0.001** |
|  |  | DietH | 0.056 | 0.101 | 0.558 | 0.577 |
|  |  | DietO | -0.111 | 0.129 | -0.865 | 0.388 |
|  |  | HabitatMarine | -0.626 | 0.471 | -1.330 | 0.184 |
|  |  | Size | -0.029 | 0.048 | -0.598 | 0.550 |
| Invertebrate | DietC HabitatFreshwater | (Intercept) | 5.271 | 0.081 | 65.022 | **< 0.001** |
|  |  | DietD | -0.187 | 0.110 | -1.697 | 0.090 |
|  |  | DietH | 0.457 | 0.072 | 6.370 | **< 0.001** |
|  |  | DietO | 0.248 | 0.092 | 2.691 | **0.007** |
|  |  | HabitatMarine | -0.111 | 0.070 | -1.578 | 0.115 |
|  |  | HabitatTerrestrial | -0.002 | 0.206 | -0.009 | 0.993 |
|  |  | Size | 0.081 | 0.019 | 4.266 | **< 0.001** |
| Mammal | DietC | (Intercept) | -8.965 | 9.520 | -0.942 | 0.383 |
|  |  | DietH | -3.093 | 2.701 | -1.145 | 0.296 |
|  |  | DietO | -4.838 | 4.092 | -1.182 | 0.282 |
|  |  | Size | 4.320 | 2.120 | 2.037 | 0.088 |
| Sauropsid | DietC HabitatFreshwater | (Intercept) | 5.249 | 0.460 | 11.407 | **< 0.001** |
|  |  | DietH | -1.226 | 0.777 | -1.578 | 0.121 |
|  |  | DietO | 0.250 | 0.264 | 0.947 | 0.348 |
|  |  | HabitatMarine | -1.070 | 1.119 | -0.956 | 0.344 |
|  |  | HabitatTerrestrial | 0.020 | 0.276 | 0.073 | 0.942 |
|  |  | Size | -0.110 | 0.232 | -0.473 | 0.638 |

**Table S9**: Model parameters for body C:P ratio as a function of diet, habitat and body size for each group of heterotrophs. *Fish1*: Euteleosteomorpha, *Fish2*: Otomorpha, *DietC*: carnivore, *DietD*: detritivore, *DietH*: herbivore, *DietO*: omnivore, *HabitatFreshwater*: freshwater, *HabitatMarine*: marine, *HabitatTerrestrial*: terrestrial, *Size*: log_10_(dry mass). P-values < 0.05 are shown in bold.

| Group | Level of comparison | Parameter | Estimate | Std. Error | t value | Pr(>\|t\|) |
| --- | --- | --- | --- | --- | --- | --- |
| Amphibian | DietC | (Intercept) | 31.746 | 8.009 | 3.964 | **< 0.001** |
|  |  | DietH | 202.470 | 11.892 | 17.025 | **< 0.001** |
|  |  | DietO | 310.035 | 20.514 | 15.113 | **< 0.001** |
|  |  | Size | -9.758 | 5.824 | -1.675 | **0.095** |
| Fish1 | Diet HabitatFreshwater | (Intercept) | 40.042 | 1.274 | 31.440 | **< 0.001** |
|  |  | DietH | 9.131 | 3.886 | 2.350 | **0.019** |
|  |  | DietO | -11.196 | 2.916 | -3.840 | **< 0.001** |
|  |  | HabitatMarine | 29.136 | 4.153 | 7.015 | **< 0.001** |
|  |  | Size | 2.299 | 1.026 | 2.242 | **0.025** |
| Fish2 | DietC HabitatFreshwater | (Intercept) | 43.385 | 1.484 | 29.233 | **< 0.001** |
|  |  | DietH | 2.635 | 1.970 | 1.338 | 0.182 |
|  |  | DietO | 2.634 | 2.510 | 1.049 | 0.295 |
|  |  | HabitatMarine | 12.458 | 9.184 | 1.356 | 0.176 |
|  |  | Size | 0.552 | 0.943 | 0.586 | 0.558 |
| Invertebrate | DietC HabitatFreshwater | (Intercept) | 238.702 | 6.435 | 37.092 | **< 0.001** |
|  |  | DietD | -69.624 | 20.205 | -3.446 | **0.001** |
|  |  | DietH | -62.801 | 5.492 | -11.436 | **< 0.001** |
|  |  | DietO | -66.613 | 6.691 | -9.956 | **< 0.001** |
|  |  | HabitatMarine | -77.719 | 5.330 | -14.580 | **< 0.001** |
|  |  | HabitatTerrestrial | -54.485 | 16.586 | -3.285 | **0.001** |
|  |  | Size | -8.913 | 1.365 | -6.531 | **< 0.001** |
| Sauropsid | DietC | (Intercept) | 12.597 | 4.582 | 2.749 | **0.010** |
|  |  | DietO | 1.880 | 1.073 | 1.753 | 0.090 |
|  |  | Size | -0.824 | 2.778 | -0.297 | 0.769 |

**Table S10**: Model parameters for body N:P ratio as a function of diet, habitat and body size for each group of heterotrophs. *Fish1*: Euteleosteomorpha, *Fish2*: Otomorpha, *DietC*: carnivore, *DietD*: detritivore, *DietH*: herbivore, *DietO*: omnivore, *HabitatFreshwater*: freshwater, *HabitatMarine*: marine, *HabitatTerrestrial*: terrestrial, *Size*: log_10_(dry mass). P-values < 0.05 are shown in bold.

| Group | Level of comparison | Parameter | Estimate | Std. Error | t value | Pr(>\|t\|) |
| --- | --- | --- | --- | --- | --- | --- |
| Amphibian | DietC HabitatFreshwater | (Intercept) | 2.822 | 0.951 | 2.966 | **0.003** |
|  |  | DietH | 20.016 | 1.418 | 14.120 | **< 0.001** |
|  |  | DietO | 16.859 | 1.499 | 11.248 | **< 0.001** |
|  |  | HabitatTerrestrial | -11.407 | 3.436 | -3.320 | **0.001** |
|  |  | Size | -1.887 | 0.686 | -2.750 | **0.006** |
| Fish1 | DietC HabitatFreshwater | (Intercept) | 3.789 | 0.090 | 42.198 | **< 0.001** |
|  |  | DietH | 0.528 | 0.281 | 1.880 | 0.061 |
|  |  | DietO | -0.855 | 0.200 | -4.276 | **< 0.001** |
|  |  | HabitatMarine | 1.382 | 0.240 | 5.769 | **< 0.001** |
|  |  | Size | 0.399 | 0.066 | 6.081 | **< 0.001** |
| Fish2 | DietC HabitatFreshwater | (Intercept) | 3.711 | 0.149 | 24.854 | **< 0.001** |
|  |  | DietH | 0.445 | 0.198 | 2.248 | **0.025** |
|  |  | DietO | 0.317 | 0.253 | 1.254 | 0.210 |
|  |  | HabitatMarine | 1.699 | 0.924 | 1.839 | 0.067 |
|  |  | Size | 0.048 | 0.095 | 0.503 | 0.615 |
| Invertebrate | DietC HabitatFreshwater | (Intercept) | 20.165 | 0.558 | 36.127 | **< 0.001** |
|  |  | DietD | -5.684 | 1.810 | -3.141 | **0.002** |
|  |  | DietH | -6.504 | 0.479 | -13.593 | **< 0.001** |
|  |  | DietO | -6.582 | 0.586 | -11.237 | **< 0.001** |
|  |  | HabitatMarine | -4.799 | 0.457 | -10.504 | **< 0.001** |
|  |  | HabitatTerrestrial | -3.718 | 1.453 | -2.558 | **0.011** |
|  |  | Size | -0.712 | 0.117 | -6.083 | **< 0.001** |
| Mammal | DietC | (Intercept) | 5.251 | 0.709 | 7.406 | **< 0.001** |
|  |  | DietH | -0.420 | 0.687 | -0.612 | 0.546 |
|  |  | DietO | 0.141 | 0.689 | 0.205 | 0.839 |
|  |  | Size | -0.343 | 0.094 | -3.670 | **0.001** |
| Sauropsid | DietC HabitatFreshwater | (Intercept) | 0.913 | 0.552 | 1.655 | 0.106 |
|  |  | DietO | 0.213 | 0.341 | 0.626 | 0.535 |
|  |  | HabitatTerrestrial | 3.129 | 0.401 | 7.810 | **< 0.001** |
|  |  | Size | 0.132 | 0.255 | 0.520 | 0.606 |

**Figure S11** Magnitude of the stoichiometric drivers for C content assessed with sequential ANCOVAs. (a) taxonomic groups, (b) diet, (c) habitat and (d) body mass. Raw residuals (points) and distributions (kernel densities) are depicted with the number of observations (at the bottom for panels a-c). Different letters indicate significant differences with post-hoc Dunn’s test of multiple comparisons with Bonferroni corrections (panels a-c). The red line panel (d) show the prediction of body C content residuals regressed against log-transformed dry body masses.

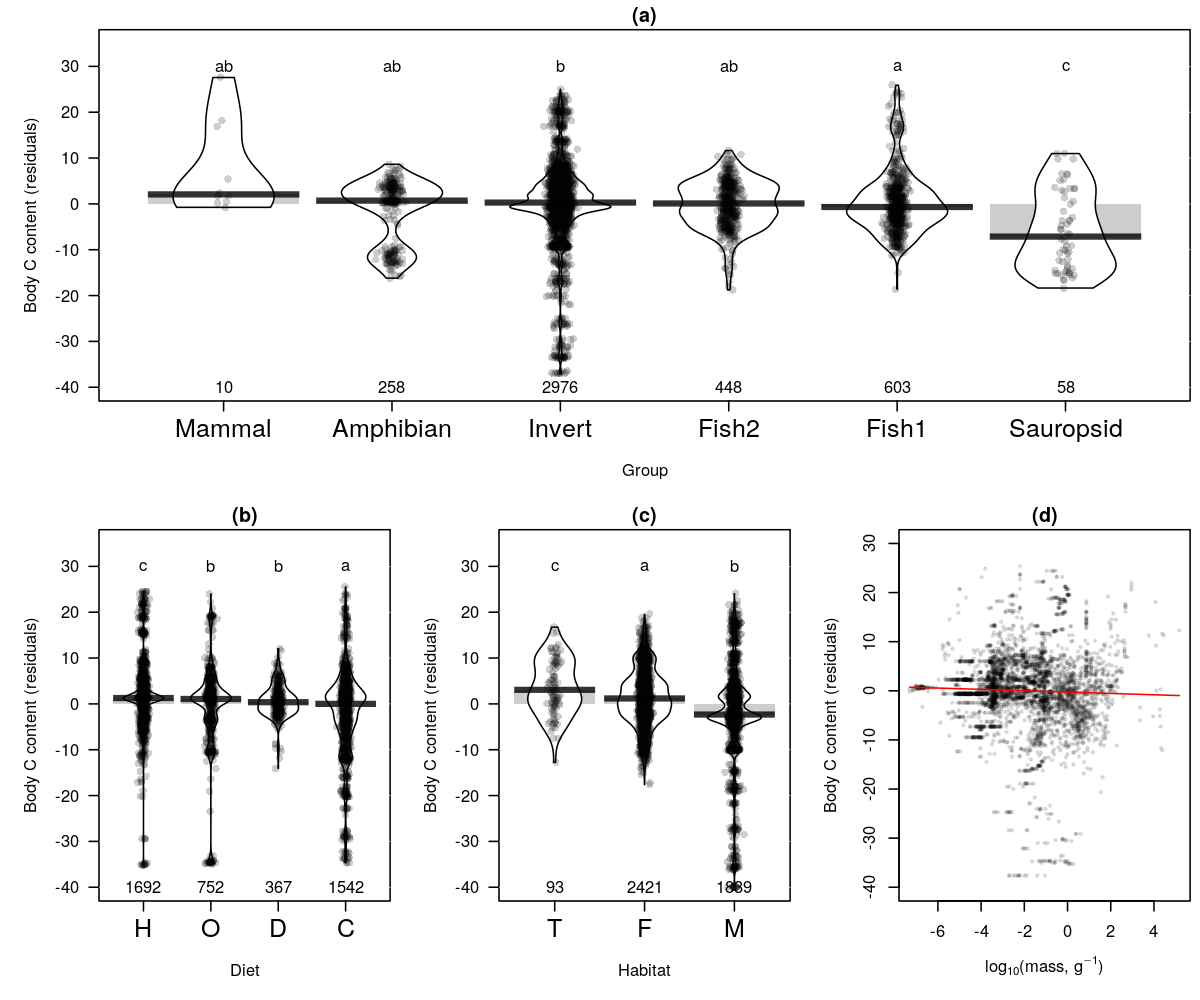

**Figure S12** Magnitude of the stoichiometric drivers for N content assessed with sequential ANCOVAs. (a) taxonomic groups, (b) diet, (c) habitat and (d) body mass. Raw residuals (points) and distributions (kernel densities) are depicted with the number of observations (at the bottom for panels a-c). Different letters indicate significant differences with post-hoc Dunn’s test of multiple comparisons with Bonferroni corrections (panels a-c). The red line panel (d) show the prediction of body N content residuals regressed against log-transformed dry body masses.

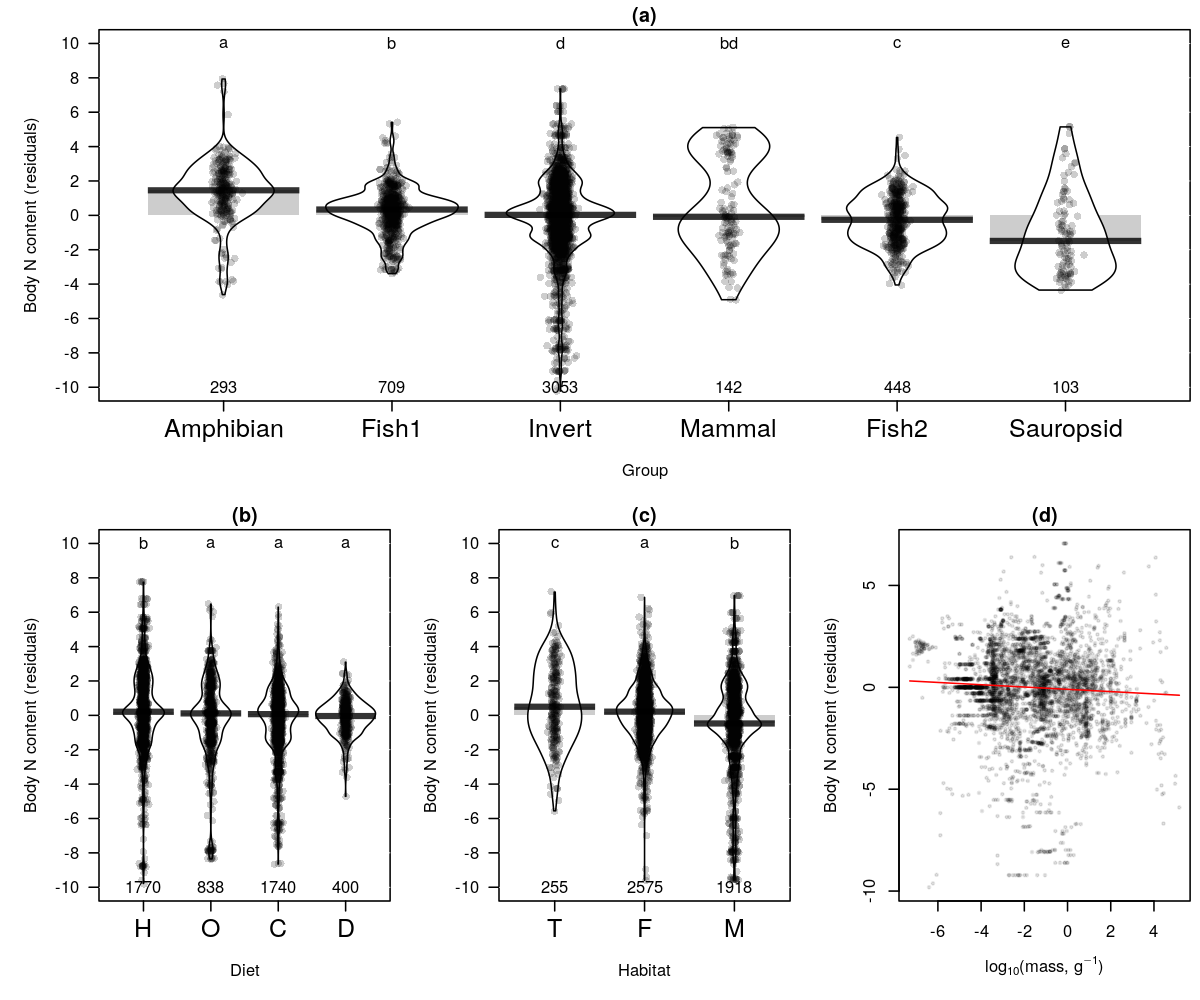

**Figure S13** Magnitude of the stoichiometric drivers for C:N assessed with sequential ANCOVAs. (a) taxonomic groups, (b) diet, (c) habitat and (d) body mass. Raw residuals (points) and distributions (kernel densities) are depicted with the number of observations (at the bottom for panels a-c). Different letters indicate significant differences with post-hoc Dunn’s test of multiple comparisons with Bonferroni corrections (panels a-c). The red line panel (d) show the prediction of body C:N residuals regressed against log-transformed dry body masses.

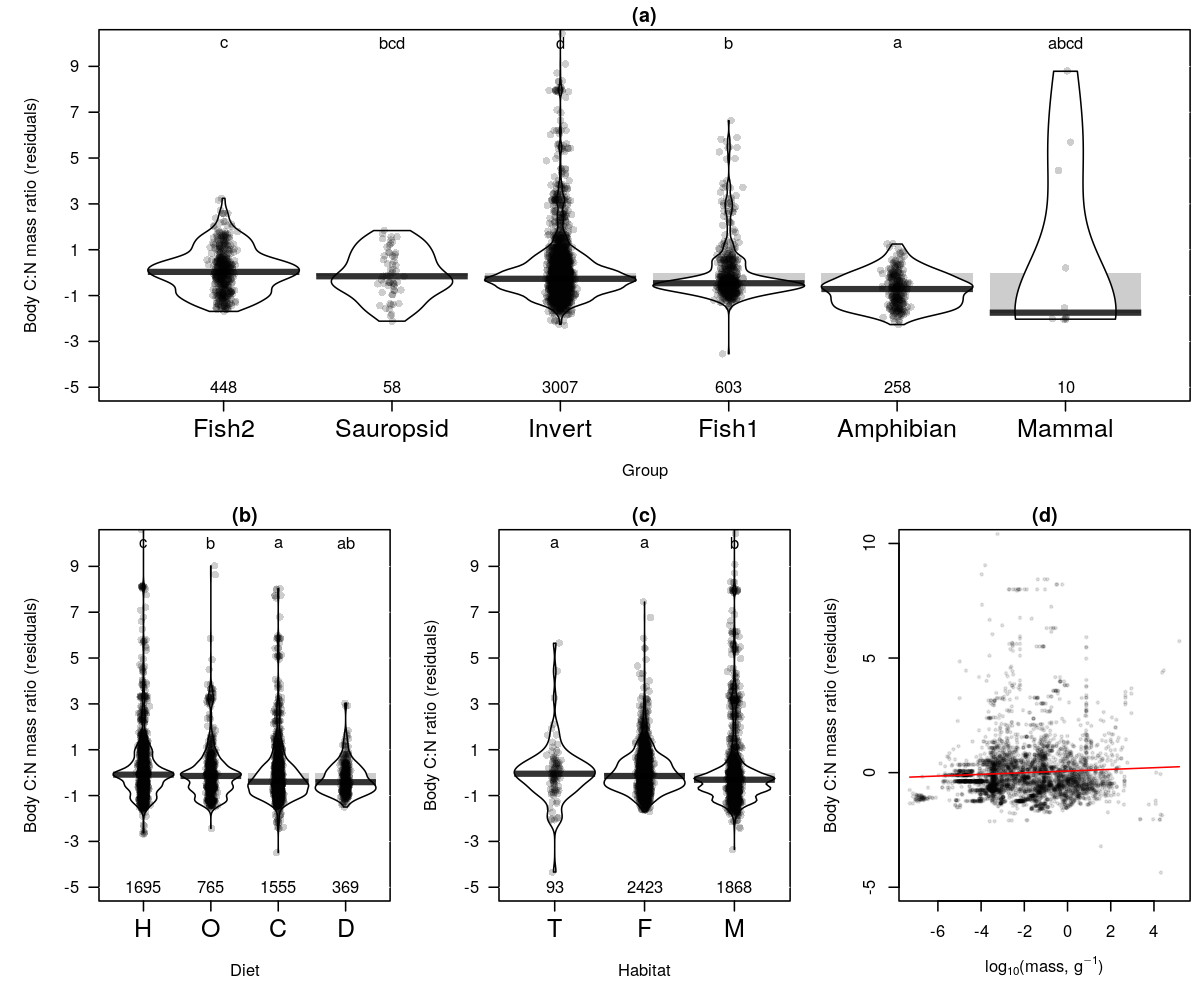

**Figure S14** Magnitude of the stoichiometric drivers for C:P assessed with sequential ANCOVAs. (a) taxonomic groups, (b) diet, (c) habitat and (d) body mass. Raw residuals (points) and distributions (kernel densities) are depicted with the number of observations (at the bottom for panels a-c). Different letters indicate significant differences with post-hoc Dunn’s test of multiple comparisons with Bonferroni corrections (panels a-c). The red line panel (d) show the prediction of body C:P residuals regressed against log-transformed dry body masses.

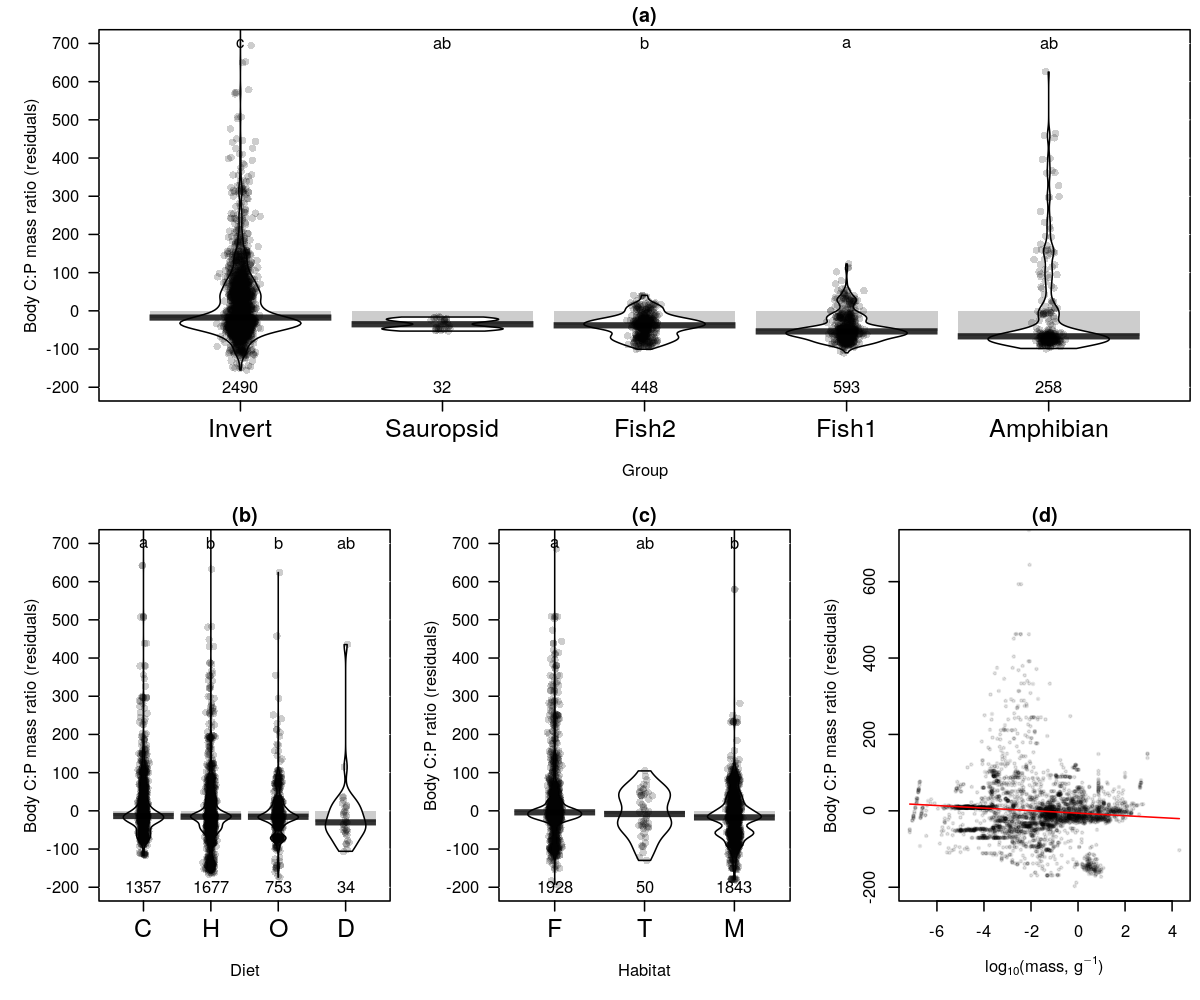

**Figure S15** Magnitude of the stoichiometric drivers for N:P assessed with sequential ANCOVAs. (a) taxonomic groups, (b) diet, (c) habitat and (d) body mass. Raw residuals (points) and distributions (kernel densities) are depicted with the number of observations (at the bottom for panels a-c). Different letters indicate significant differences with post-hoc Dunn’s test of multiple comparisons with Bonferroni corrections (panels a-c). The red line panel (d) show the prediction of body N:P residuals regressed against log-transformed dry body masses.

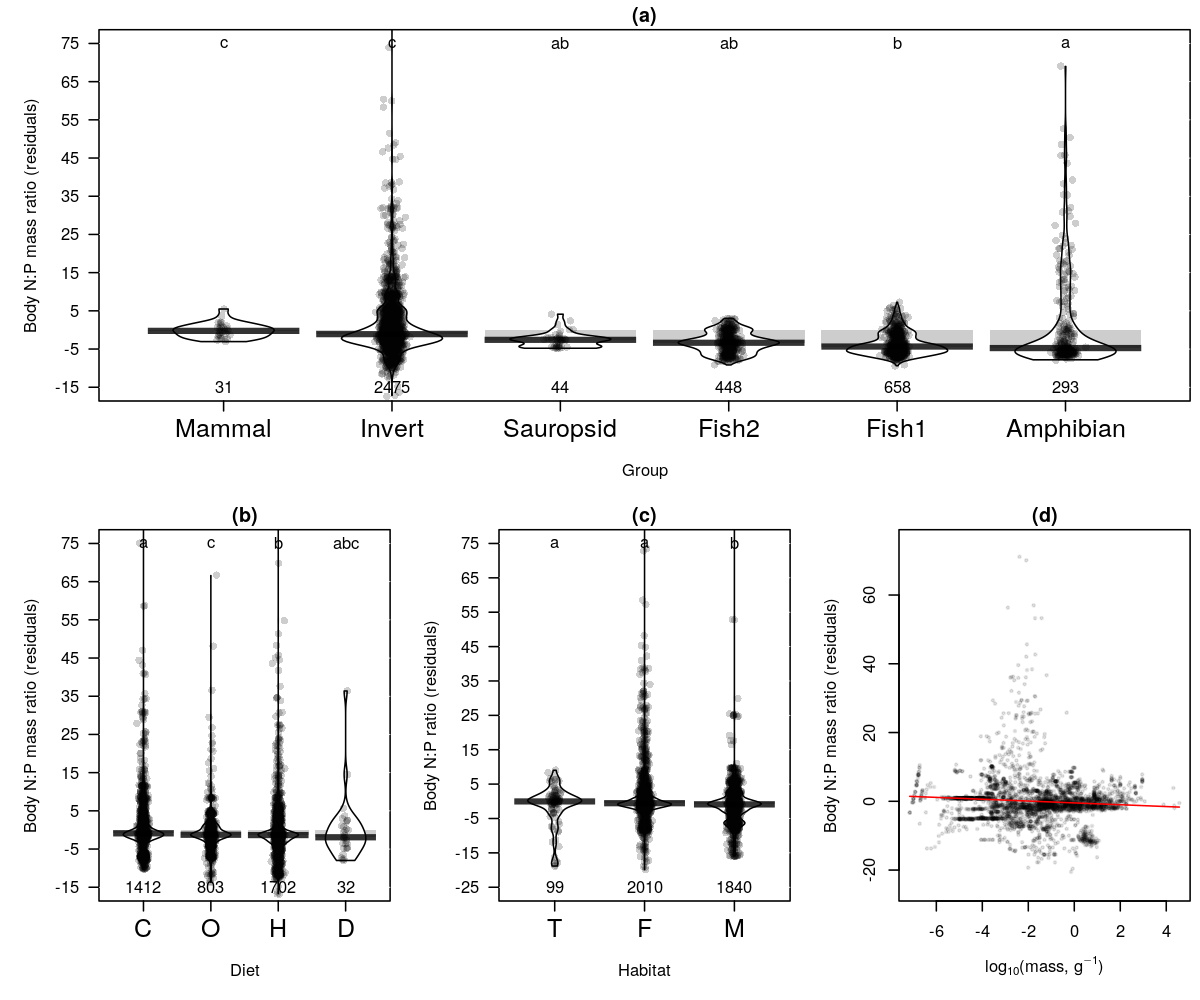

**Table S16**: Phylogenetic variation of the six stoichiometric traits analyzed both with Blomberg’s K- and Pagel’s lambda- statistics.

| Stoichiometric trait | n | Blomberg's K | p-value | Pagel's lambda | LL | LL0 | p-value |
| --- | --- | --- | --- | --- | --- | --- | --- |
| C | 163 | 0.071 | 0.010 | 0.920 | -477.575 | -499.723 | < 0.001 |
| N | 273 | 0.379 | 0.001 | 0.978 | -515.027 | -655.370 | < 0.001 |
| P | 221 | 0.500 | 0.001 | 0.958 | -222.739 | -333.487 | < 0.001 |
| C:N | 174 | 0.493 | 0.012 | 0.996 | -494.803 | -597.977 | < 0.001 |
| C:P | 161 | 0.707 | 0.001 | 1.000 | -871.905 | -981.787 | < 0.001 |
| N:P | 173 | 0.718 | 0.001 | 1.000 | -450.592 | -563.042 | < 0.001 |

LL: log-likelihood, LL0: log-likelyhood for lambda = 0.0.
